# Cyst-type epithelial heterogeneity shapes therapeutic responsiveness in ADPKD

**DOI:** 10.64898/2026.05.08.723838

**Authors:** Johannes Roeles, Axel Giesler, Jessica Schmitz, Johannes König, Frank Friedersdorff, Jonas Busch, Robert Greite, Matteus Krappitz, Isabel A. Layne, Michael Köttgen, E. Wolfgang Kuehn, Jessica Schmid, Katharina Schmidt, Konrad Hoeft, Rafael Kramann, Kai-Uwe Eckardt, Christine Kocks, Jan H. Bräsen, Jan Halbritter, Roland Schmitt, Silke Härteis, Katharina Hopp, Kai M. Schmidt-Ott, Christian Hinze

## Abstract

Autosomal dominant polycystic kidney disease (ADPKD) exhibits substantial interpatient variability in disease course and therapeutic response, but the cellular basis for this variability remains poorly understood. Here, we combine single-nucleus RNA sequencing of human cyst epithelia with machine learning–based histological analysis of >1,800 cysts to resolve three epithelial cyst types—proximal tubule–like, collecting duct–like, and mixed. These cyst types display distinct injury states, metabolic programs, and stromal microenvironments, including a mixed-cyst niche enriched for CCL2-associated inflammatory signaling. Expression of key therapeutic targets was highly cell-type specific with CFTR enriched in proximal-like epithelia, whereas AVPR2 expression was confined to AQP2-positive collecting duct–like cells. Cyst-type composition varied widely across patients and in an orthologous mouse model (*Pkd1*^RC/RC^) in which the burden of AQP2-positive cysts correlated with responsiveness to tolvaptan. These findings identify cyst-type heterogeneity as a major determinant of molecular pathway activation and predictability of therapeutic response in ADPKD.

## Introduction

Autosomal dominant polycystic kidney disease (ADPKD) is the most common inherited kidney disease, affecting at least 1 in 1,000 individuals (*1, 2*). It is characterized by progressive cystic remodeling of the kidney that culminates in end-stage kidney disease and renal replacement therapy (*3*). More than 90% of cases are due to mutations to *PKD1* or *PKD2*, which encode polycystin 1 and polycystin 2, transmembrane proteins that form heterooligomeric complexes in the primary cilium and other compartments (*3*). Although the polycystin complex is thought to help regulate tubule caliber and resist cystic dilation (*4*), the mechanisms of disease initiation and progression remain incompletely defined. Moreover, clinical heterogeneity is pronounced: patients with *PKD1* mutations typically progress faster than those with *PKD2*, yet severity varies widely even among relatives with the same primary mutation (*5*). Tolvaptan, an arginine vasopressin receptor 2 (AVPR2) antagonist and the only approved therapy for ADPKD, shows variable effectiveness across patients and can cause polyuria and liver toxicity (*6*), limiting broad applicability and long-term tolerance. In view of the urgent need for novel treatment options, numerous therapeutic strategies are under preclinical and clinical evaluation, including agents targeting metabolic reprogramming, epithelial proliferation, immune-mediated injury, oxidative stress, and abnormal solute-water transport (*7*). However, translation to consistent patient benefit remains challenging. One barrier to translation is limited understanding of molecular heterogeneity within human ADPKD cysts. To address this, we combined single-nucleus RNA sequencing (snRNA-seq) with machine learning-aided image analysis to characterize intra- and interpatient cyst-type diversity. SnRNA-seq resolved three cyst types with different epithelial layers – proximal tubule-like, collecting duct-like, and mixed cysts – each characterized by distinct injury states, metabolic programs, and stromal microenvironments. Distinct expression of molecular drug targets indicated differential molecular targetability across cyst types and machine-learning–guided tissue profiling uncovered substantial interpatient variability in cyst-type composition, with CD-like to PT-like ratios ranging from 1:3.5 to 8:1 across kidneys. Finally, in the *Pkd1*^RC/RC^ mouse model, tolvaptan treatment response tracked with the burden of AQP2^+^ cysts, supporting a mechanistic link between cyst-type composition and responsiveness to AVPR2 antagonism. Together, these findings delineate previously underappreciated cyst-type heterogeneity in human ADPKD and suggest that epithelial lineage and cyst-type composition may contribute to interpatient variability in disease progression and treatment response.

## Results

### Human ADPKD cyst epithelia exhibit pure or mixed cellular identities

We collected 15 superficial cyst wall samples from five late-stage ADPKD nephrectomies and tumor-adjacent normal kidney tissue from three patients undergoing nephrectomy for renal tumors (Fig. 1A; Suppl. Table 1). All cysts were profiled by single-nucleus RNA sequencing (snRNA-seq; Chromium Next GEM Single Cell 3′ v3.1) and microscopy, and cyst fluid was sampled from each sequenced cyst. Quality control and downstream analyses were performed in Seurat (*8*) with dataset integration and batch correction using Harmony (*9*); cell types were annotated prior to integration to avoid batch-driven misassignment. Three cyst samples did not meet sequencing quality thresholds due to low epithelial cell recovery and were excluded from quantitative analyses. The integrated dataset of ADPKD and control tissue recovered all major kidney cell types (Fig. 1B; Suppl. Fig. 3). Consistent with prior reports, cyst samples contained a higher fraction of fibroblasts and immune cells than normal tissue (Fig. 1C; Suppl. Fig. 4).

**Figure 1.**
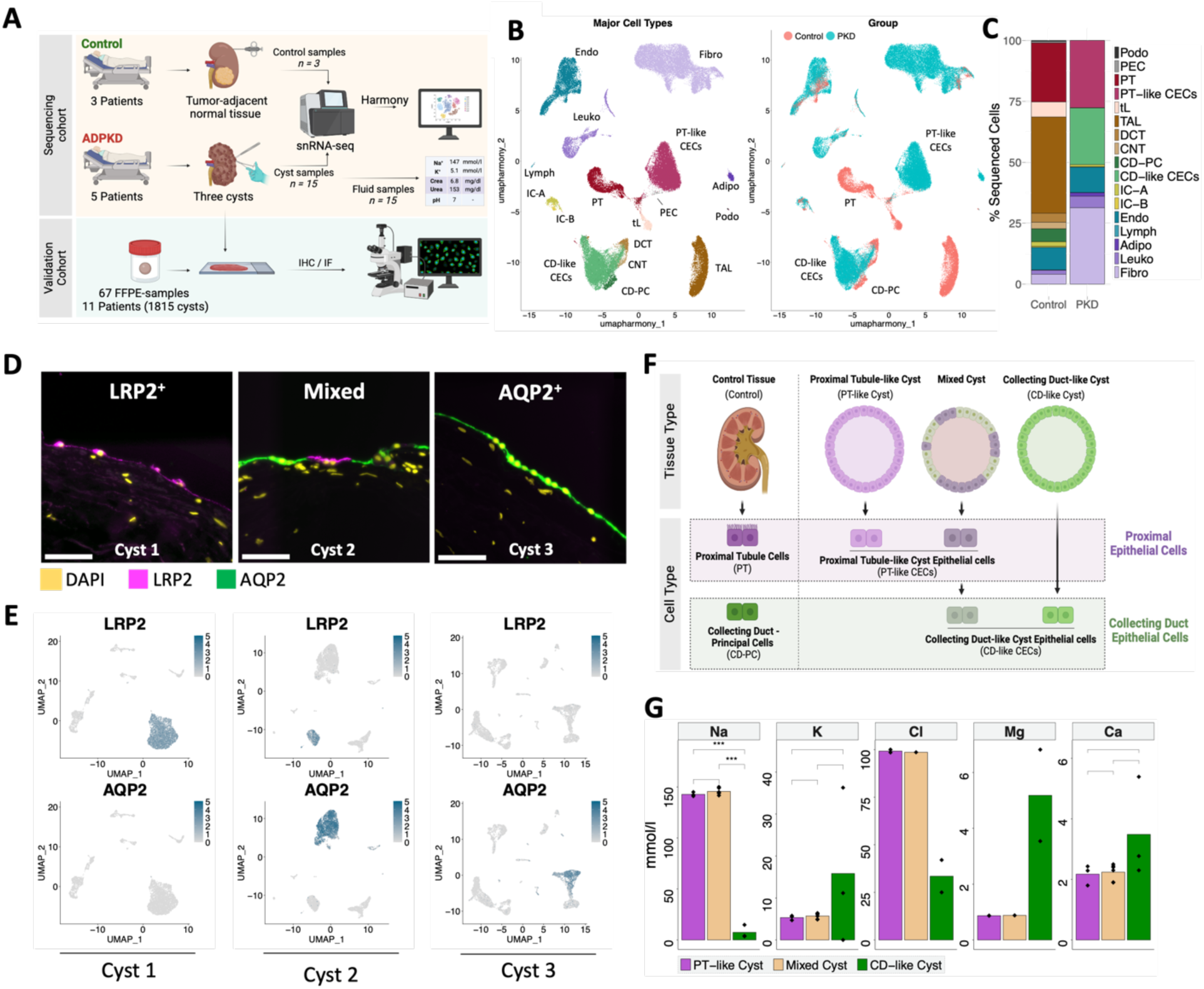
ADPKD cyst epithelia exhibit pure or mixed cellular identities. (**A**) Schematic of the study design. (**B**) Uniform manifold approximation and projection (UMAP) of the integrated Seurat object. Left, major kidney cell types; right, tissue of origin (ADPKD vs control). (**C**) Fraction of major kidney cell types relative to total sequenced cells in ADPKD and control samples. (**D**) Representative immunofluorescence images showing LRP2 (magenta) and AQP2 (green) in different cysts. Nuclei are stained with DAPI (yellow). Scale bars, 50 µm. (**E**) Representative feature plots showing LRP2 and AQP2 expression in snRNA-seq datasets of three individual cysts. (**F**) Schematic of tissue and cell types defined in the study. (**G**) Mean concentrations of sodium (Na), potassium (K), chloride (Cl), magnesium (Mg), and calcium (Ca) by cyst type. Adipo, adipocytes; CNT, connecting tubule; DCT, distal convoluted tubule; Endo, endothelial cells; FFPE, formalin-fixed paraffin-embedded; Fibro, fibroblasts; IC-A, intercalated cells type A; IC-B, intercalated cells type B; Leuko, leukocytes; Lymph, lymphatic endothelial cells; PEC, parietal epithelial cells; Podo, podocytes; tL, thin limb. *, P ≤ 0.05; **, P ≤ 0.01; ***, P ≤ 0.001; ****, P ≤ 0.0001; no asterisk, P > 0.05.

Among tubular epithelial cells, cyst walls were lined by two lineages: proximal tubule–like cyst epithelial cells (PT-like CECs) with PT-associated gene signatures, including the conserved marker low-density lipoprotein receptor-related protein 2 (LRP2; megalin) and collecting duct-like CECs (CD-like CECs) with principal cell (CD-PC)-like signatures, including the conserved marker aquaporin-2 (AQP2). (Fig. 1C, Suppl. Fig. 3). Epithelial populations from other segments (e.g., thick ascending limb; TAL), which constitute a substantial fraction in healthy tissue, were not detected in our ADPKD cyst walls (Fig. 1C), consistent with reports that LRP2 and AQP2 are the most commonly detectable cyst markers in the *Pkd1*^RC/RC^ mouse model, which produces progressive, adult-onset polycystic kidney disease and is widely used for chronic intervention studies because it recapitulates key features of human ADPKD (*10–12*).

In a subset of cysts, PT-like and CD-like CECs coexisted within the same wall (Fig. 1D–E). Immunofluorescence revealed arrangements ranging from cell-type patches to cell-by-cell mosaic patterns and single interposed cells (Suppl. Fig. 5), with occasional LRP2/AQP2 double-positive cells. Based on these features, we defined three human ADPKD cyst types in the snRNA-seq dataset: PT-like cysts containing PT-like CECs, CD-like cysts containing CD-like CECs, and mixed cysts containing both lineages (Fig. 1F).

Cyst fluid composition tracked with epithelial identity (Fig. 1G). PT-like and mixed cyst fluids contained sodium, chloride, magnesium, calcium, and potassium concentrations similar to serum levels, whereas CD-like cysts exhibited markedly low sodium concentrations – nearly 20-fold lower than in PT-like or mixed cysts – with variable potassium levels. These patterns concord with early cyst puncture studies reporting distinct populations of high-sodium and low-sodium cysts (*13*) and support a model in which cyst-lining cell identity shapes fluid chemistry.

### Cyst epithelial cells exhibit cellular injury and metabolic alteration

To assess cyst type–specific epithelial states, we stratified CECs by lineage (PT-like vs. CD-like) and by cyst-type of origin (PT-like, CD-like, mixed), yielding four categories: PT-like CECs from PT-like cysts, PT-like CECs from mixed cysts, CD-like CECs from CD-like cysts, and CD-like CECs from mixed cysts (Fig. 1F). Subclustering with control PT and CD-PC cells revealed that PT-like CECs formed two clusters (Clust1, Clust2) distinct from healthy PT and vascular cell adhesion molecule 1 (*VCAM1*)-positive failed-repair (*14*) PT cells (Fig. 2A; Suppl. Fig. 6). Clust2 was enriched for PT-like CECs from PT-like cysts, whereas Clust1 comprised predominantly PT-like CECs from mixed cysts (Fig. 2B).

**Figure 2.**
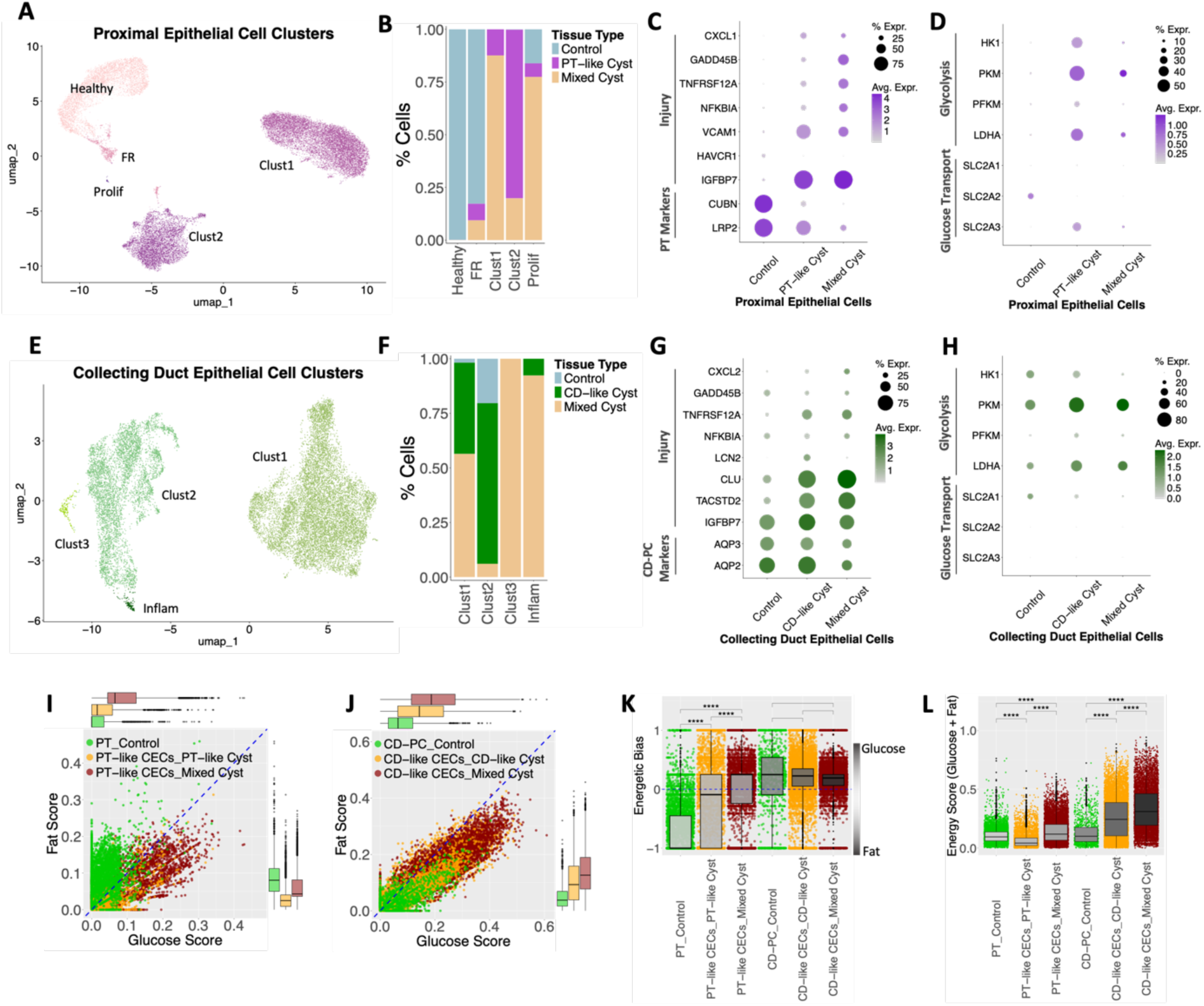
Cyst epithelial cell types display injury and metabolic alterations. (**A**) UMAP of proximal epithelial cells (PT-like CECs and control PT cells). (**B**) Tissue-type representation across proximal epithelial clusters. (**C**) Dot plot showing expression of injury-associated genes in proximal epithelial cells by tissue type. (**D**) Dot plot showing expression of key genes involved in glucose transport and glycolysis in proximal epithelial cells by tissue type. (**E**) UMAP of collecting duct epithelial cells (CD-like CECs and control CD-PCs). (**F**) Tissue-type representation across collecting duct epithelial clusters. (**G**) Dot plot showing expression of injury- and ECM-associated genes in collecting duct epithelial cells by tissue type. (**H**) Dot plot showing expression of key genes involved in glucose transport and glycolysis in collecting duct epithelial cells by tissue type. (**I–J**) Scatter plots with adjacent box plots showing expression scores of energy metabolism-associated genes by cell type and tissue type. The dashed blue line indicates equal glucose and fat scores. (**K**) Scatter plot with adjacent box plot showing energetic bias (glucose-to-fat score ratio) by cell type and tissue type. (**L**) Scatter plot with adjacent box plots showing combined energy metabolism score by cell type and tissue type. FR, failed-repair proximal tubule cells; Healthy, healthy proximal tubule cells; Prolif, proliferating cells; *SLC2A1*, solute carrier family 2 member 1 (GLUT1). *, P ≤ 0.05; **, P ≤ 0.01; ***, P ≤ 0.001; ****, P ≤ 0.0001; no asterisk, P > 0.05.

Across PT-like CECs, injury and epithelial–mesenchymal transition (EMT) markers, such as *VCAM1* and insulin-like growth factor-binding protein 7 (*IGFBP7*), were elevated, with the strongest signals in cells from mixed cysts (Fig. 2C). PT-like CECs from mixed cysts also showed upregulation of nuclear factor kappa B (NF-κB) pathway genes (e.g., NF-κB inhibitor alpha, *NFKBIA*) and tumor necrosis factor alpha (TNF-α) signaling components (e.g., tumor necrosis factor receptor superfamily member 12A, *TNFRSF12A*), consistent with reports that *TNFRSF12A* axis promotes inflammatory responses in the cyst epithelium and microenvironment (*15*). Expression of growth arrest and DNA-damage-inducible beta (*GADD45B*) and C-X-C motif chemokine ligand 1 (*CXCL1*) was also increased. Gene-level changes were corroborated by pathway scoring, with increased EMT and TNF-α activity in PT-like CECs from mixed cysts (Suppl. Fig. 6C). These injury programs coincided with reduced expression of canonical PT markers such as *LRP2* and and cubilin (*CUBN*), indicating progressive dedifferentiation of PT-like CECs (Fig. 2C).

The common kidney injury marker hepatitis A virus cellular receptor 1 (*HAVCR1*) was not expressed PT-like CECs, indicating a state of cell injury in cyst epithelial cells that is distinct from non-cystic tubular epithelial cells (Suppl. Fig 6B).

Along with dedifferentiation and injury, core metabolic programs were altered. Key glycolysis genes were upregulated in PT-like CECs relative to control PT cells, including hexokinase 1 (*HK1*), phosphofructokinase, muscle (*PFKM*), pyruvate kinase M1/2 (*PKM*), and lactate dehydrogenase A (*LDHA*) (Fig. 2D). Conversely, genes involved in fatty acid metabolism – the primary energy source in healthy proximal tubules – were reduced in PT-like CECs (Suppl. Fig. 7), consistent with a glycolytic shift in LRP2^+^ cyst epithelia previously reported in ADPKD (*16, 17*). Pathways mediating gluconeogenesis and transcellular reabsorption of glucose and amino acids were markedly diminished in PT-like CECs, especially those from mixed cysts (Suppl. Fig. 8). This included reduced expression of sodium/glucose cotransporter 2 (*SLC5A2*; SGLT2) and glucose transporter type 2 (*SLC2A2*; GLUT2), which in healthy PT facilitate apical uptake of filtered glucose and basolateral release, respectively (*18*). Instead, glucose transporter type 3 (*SLC2A3*; GLUT3) was expressed, which is primarily expressed in neurons and warrants intracellular glucose supply for energy production even at low blood glucose concentrations (*19*).

Subclustering of collecting duct lineage cells yielded four distinct clusters. Control CD-PC were almost exclusively found in Clust2 while Clust1 comprised CD-like CECs from, both, CD-like and mixed cysts. Clust3 and an inflammatory cluster (Inflam) were predominated by mixed cyst cells. CD-like CECs from pure vs. mixed cysts exhibited a related pattern of injury as PT-like CECs while genes involved in glucose metabolism were not markedly changed (Fig. 2E–H; Suppl. Fig. 6). Collecting duct injury markers such as clusterin (*CLU*) and tumor-associated calcium signal transducer 2 (*TACSTD2*) were upregulated, along with components of tumor necrosis factor alpha (TNF-α) signaling (*TNFRSF12A*) and chemotaxis pathways (C-X-C motif chemokine ligand 2, *CXCL2*). The kidney injury marker lipocalin-2 (*LCN2*) was detected only in CD-like CECs from CD-like cysts, but not from mixed cysts.

To quantify metabolic differences across cyst types, we computed single-cell gene expression scores for core glucose and fatty acid metabolism. The glucose score summarized genes involved in glucose uptake and glycolysis; the fat score captured genes driving fatty acid uptake, mitochondrial import, and beta-oxidation. Genes involved in oxidative phosphorylation were included in both scores (Methods). Energetic bias was defined as the ratio of the glucose to fat scores. In PT-like CECs, metabolism shifted toward glucose dependency in a cyst type–dependent manner: PT-like CECs from mixed cysts displayed higher glucose scores and greater energetic bias towards glucose metabolism than PT-like CECs from PT-like cysts (Fig. 2I, K). In contrast, CD-like CECs showed stable global substrate preference, reflected by a constant energetic bias (Fig. 2J, K). However, the total energy metabolism score – combining glucose and fatty acid pathways – was significantly higher in CD-like CECs than in PT-like CECs (Fig. 2L).

Together, these data indicate that cyst-lining epithelia adopt distinct injury states and metabolic programs in ADPKD. Injury and metabolic reprogramming increase glucose dependency, especially in PT-like CECs from mixed cysts, while CD-like CECs exhibit the highest inferred energy demand by gene expression scoring.

### ADPKD cyst epithelia are embedded in distinct cellular microenvironments

Multiple cellular compartments contribute to chronic kidney disease progression, including immune cells, fibroblasts, and endothelial cells (EC) (*20–22*). To examine how these cells may influence epithelial injury and cyst growth in human ADPKD, we analyzed single-cyst microenvironments enabled by the sampling of superficial cyst cups (Fig. 1A), which minimized confounding effects from adjacent cysts (Fig. 3; Suppl. Fig. 9).

**Figure 3.**
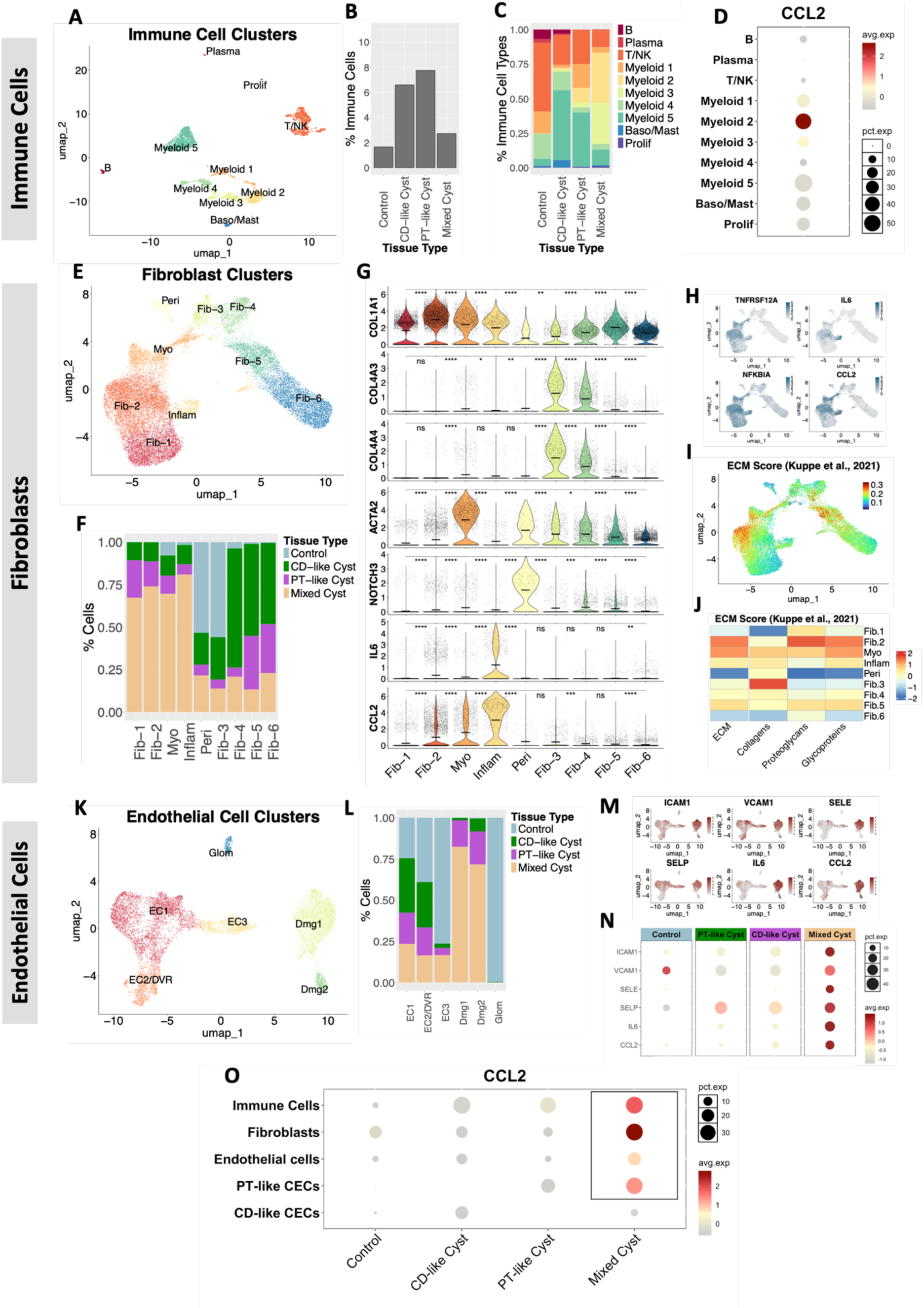
Different cyst types are embedded within distinct inflammatory microenvironments. Subclustering analyses of immune cells (**A-D**), fibroblasts (**E-J**), and endothelial cells (**K-O**). (**A**) UMAP of immune cell clusters. (**B**) Fraction of immune cells among all sequenced cells by tissue type. (**C**) Representation of immune cell clusters by tissue type. (**D**) Dot plot showing *CCL2* expression across immune clusters. (**E**) UMAP of fibroblast clusters. (F) Tissue-type representation across fibroblast clusters. (**G**) Violin plots showing expression of selected genes in fibroblast subclusters. Horizontal lines indicate mean expression. (**H**) Feature plots showing expression of selected injury markers in fibroblasts. (**I**) Feature plot showing expression of ECM genes. (**J**) Heat map of expression scores for total ECM, collagens, proteoglycans, and glycoproteins. Gene sets for (**I–J**) were obtained from Kuppe et al. (*29*). Scores were calculated using UCell (*30*). (**K**) UMAP of EC clusters. (**L**) Tissue-type representation across EC clusters. (**M-N**) Feature and dot plots showing expression of selected EC activation genes. (**O**) Dot plots showing *CCL2* expression by cell type and tissue type. *ACTA2*, actin alpha 2; B, B cells; Baso/Mast, basophil granulocytes/mast cells; *COL1A1*, collagen type I alpha 1 chain; *COL4A3*, collagen type IV alpha 3 chain; *COL4A4*, collagen type IV alpha 4 chain; DVR, descending vasa recta; Fib, fibroblasts; Glom, glomerular endothelial cells; *NOTCH3*, notch receptor 3; Peri, pericytes; Plasma, plasma cells. *, P ≤ 0.05; **, P ≤ 0.01; ***, P ≤ 0.001; ****, P ≤ 0.0001; no asterisk, P > 0.05.

In the immune compartment of control kidney tissue, T lymphocytes and natural killer (T/NK) cells predominated, whereas cyst-associated immune populations were mainly myeloid cells (Fig. 3A–C). Myeloid cell composition varied by cyst type, with marked expansion of the clusters myeloid 2 and myeloid 3 around mixed cysts (Fig. 3C), and myeloid 5 around PT-like and CD-like cysts. All three clusters expressed cluster of differentiation 14 (*CD14*) but not integrin subunit alpha X (*ITGAX*), consistent with macrophage identity (Suppl. Fig. 9A). Myeloid 3 comprised immune-active macrophages with strong expression of major histocompatibility complex (MHC) class II molecules and interleukin-1 beta (*IL1B*) (Suppl. Fig. 9). Myeloid 2 and myeloid 5 showed an M2-like macrophage phenotype akin to tumor-associated macrophages (TAMs) (*23*), marked by expression of hyaluronan mediated motility receptor (*CD168*), mannose receptor c-type 1 (*MRC1*) and folate receptor beta (*FOLR2*) (Suppl. Fig. 9B). TAMs promote tubular proliferation after injury (*24*) and have been implicated in ADPKD cyst expansion (*25, 26*). Myeloid 2 also expressed high levels of C-C motif chemokine ligand 2 (*CCL2*) (Fig. 3D), a chemotactic factor that recruits monocytes and other immune cells to sites of injury (*27*) and promotes disease progression in ADPKD models (*25, 28*).

We next profiled fibroblasts, which formed a thick layer around cysts and accounted for approximately one third of sequenced cells in ADPKD samples, irrespective of cyst type (Suppl. Fig. 10). Subclustering yielded ten fibroblast subpopulations with diverging injury-marker expression (Fig. 3E-H). Control cells were enriched in the pericyte cluster and in the collagen IV–expressing clusters Fib-3 Fibroblasts from CD-like and PT-like cysts were enriched in less-injured clusters Fib-5 and Fib-6 whereas those from mixed cysts primarily congregated in injured clusters Fib-1, Fib-2, in a distinct cluster of proinflammatory fibroblasts (Inflam), and in the myofibroblast cluster (Myo). Analysis of extracellular matrix (ECM) programs using a recent fibrosis-related gene set (*29*) showed highest ECM-associated expression in Fib-2, Fib-5, and the myofibroblast cluster (Fig. 3I–J) – consistent with the central role of myofibroblasts in ECM deposition (*29*). The cluster of proinflammatory fibroblasts was composed almost exclusively of cells from mixed cysts and characterized by high interleukin-6 (*IL6*) and *CCL2* expression (Fig. 3F, H).

Damaged EC clusters (Dmg1/2) mirrored this pattern showing elevated *IL6* and *CCL2* together with upregulation of EC activation markers, such as *VCAM1*, intercellular adhesion molecule 1 (*ICAM1*), P-selectin (*SELP*), and E-selectin (*SELE*) (Fig. 3K–N). In acute kidney injury, fibroblasts and ECs recruit leukocytes via *CCL2* signaling, and failed-repair proximal tubule cells later sustain *CCL2*-mediated chemoattraction (*14*). Consistent with this paradigm, PT-like CECs from mixed cysts exhibited highly elevated *CCL2* expression, whereas CD-like CECs showed minimal engagement in *CCL2* signaling (Fig. 3O).

Together, these analyses indicate that PT-like CECs from mixed cysts interact with specific immune, fibroblast, and endothelial subpopulations to establish a distinct inflammatory microenvironment around mixed cysts. A unifying feature of this mixed-cyst niche is high *CCL2* expression across epithelial and stromal compartments.

### Expression of molecular drug targets in CECs differs by cell type and cyst type

Given the pronounced transcriptomic differences across CECs, including metabolic programs, chemoattraction, and injury, we next asked how cell type and cyst type influence molecular amenability to therapy. We focused on pathways targeted by agents in preclinical or clinical development, including oxidative stress responses, cyclic AMP (cAMP) signaling, and solute–water transport (Fig. 4A).

**Figure 4.**
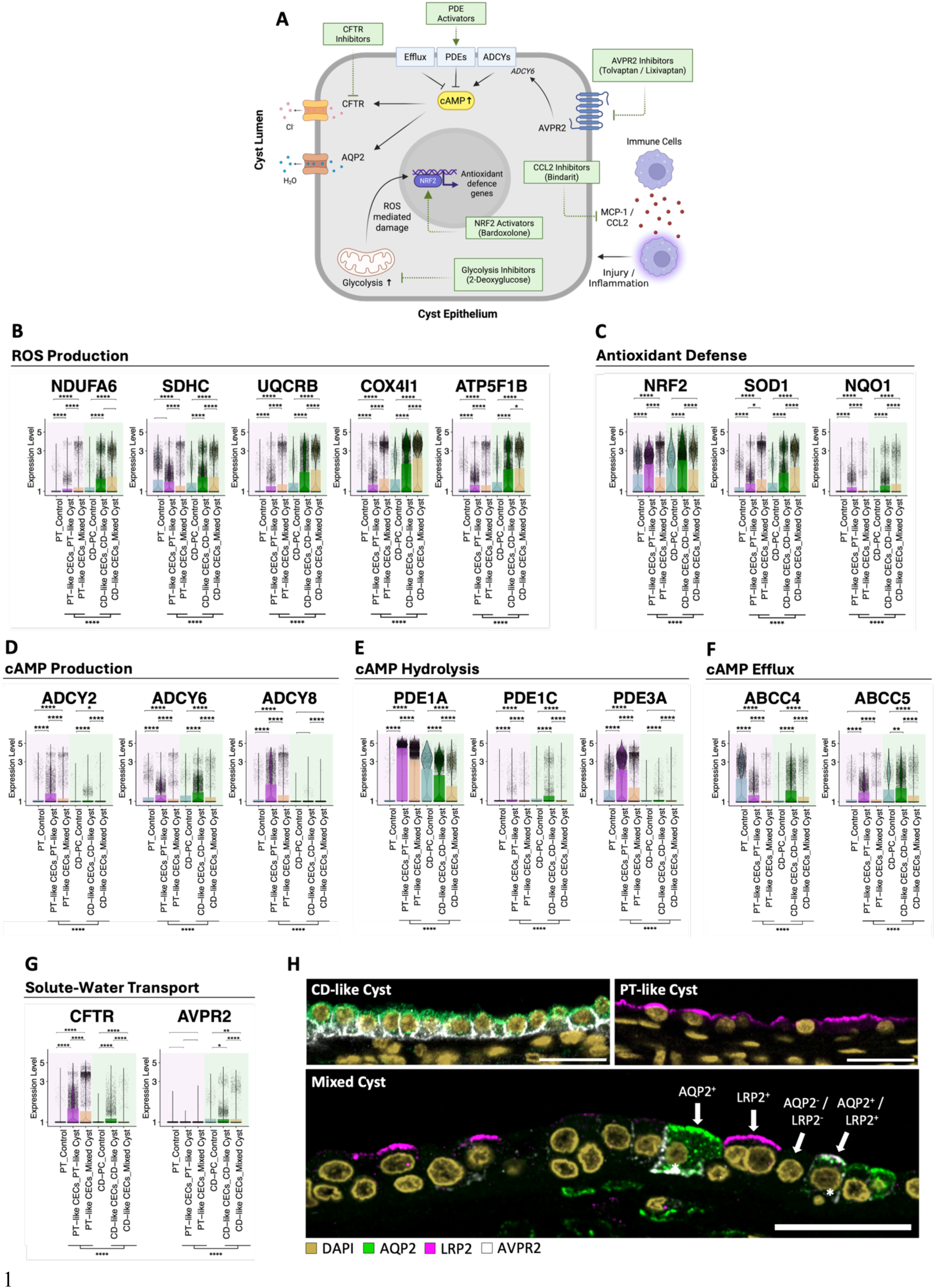
Expression of molecular drug targets varies among cyst epithelial cell types. (**A**) Schematic of selected therapeutic approaches under preclinical and clinical evaluation. (**B-F**) Violin plots showing expression of core subunits of the five mitochondrial respiratory chain complexes (**B**), core antioxidant response genes (**C**), selected adenylyl cyclases (**D**), phosphodiesterases (**E**), and cAMP efflux transporters (**F**) by cell type and cyst type; bar plots are overlaid to indicate mean expression. (**G**) Violin and feature plots showing expression of *LRP2*, *CFTR*, *AQP2*, and *AVPR2*; bars are overlaid to indicate mean expression. (**H**) Representative immunofluorescence images showing AVPR2 protein in epithelium of CD-like, PT-like, and mixed cysts. Asterisks mark cells cells expressing AVPR2 in mixed cysts. Scale bars, 30 µm. *ABCC5*, ATP-binding cassette subfamily C member 5; *ATP5F1B*, ATP synthase F1 subunit beta; *COX4I1*, cytochrome c oxidase subunit 4I1; *NDUFA6*, NADH:ubiquinone oxidoreductase subunit A6; *NQO1*, NAD(P)H quinone dehydrogenase 1; *PDE1C*, phosphodiesterase 1C; *PDE3A*, phosphodiesterase 3A; *SDHC*, succinate dehydrogenase complex subunit C; *SOD1*, superoxide dismutase 1; *UQCRB*, ubiquinol-cytochrome c reductase binding protein. *, P ≤ 0.05; **, P ≤ 0.01; ***, P ≤ 0.001; ****, P ≤ 0.0001; no asterisk, P > 0.05.

Oxidative stress is implicated in ADPKD progression (*31*) and arises from imbalances between mitochondrial reactive oxygen species (ROS) production and detoxification. Expression of core subunits of the five mitochondrial respiratory chain complexes – major cellular sources of ROS (*32*) – was markedly increased in CD-like CECs, particularly those from mixed cysts (Fig. 4B; Suppl. Fig. 11). The antioxidant regulator nuclear factor erythroid 2–related factor 2 (*NRF2*) and downstream targets, including superoxide dismutase 1 (*SOD1*) and NAD(P)H quinone dehydrogenase 1 (*NQO1*), were also upregulated in CD-like CECs (Fig. 4C). These patterns indicate a high oxidative stress burden in CD-like CECs and suggest greater susceptibility to ROS-reduction strategies.

To evaluate cAMP homeostasis, we examined expression of adenylyl cyclases (*ADCYs*), phosphodiesterases (*PDEs*), and cAMP efflux transporters, which together govern intracellular cAMP levels (*33*). *ADCY2* and *ADCY8* were upregulated in PT-like CECs but not in CD-like CECs, and *ADCY6* was elevated predominantly in CECs from pure cysts (Fig. 4D). Because ADCY6 mediates arginine vasopressin receptor 2 (AVPR2)-stimulated cAMP production (*34*), its expression may modulate responsiveness to AVPR2 antagonism. *PDE* isoforms also showed cell type-specific expression patterns, including differences in *PDE1A*, as did cAMP efflux transporters, such as multidrug resistance-associated protein 4 (*ABCC4*; MRP4) (Fig. 4E–F). Collectively, these data indicate that cAMP regulation is differentially perturbed across cyst epithelia, offering cell type–specific opportunities to modulate cAMP. Notably, selective PDE inhibitors have been reported to prevent cyst formation in vitro (*35*).

We next assessed cAMP-linked effectors implicated in ADPKD. The cystic fibrosis transmembrane conductance regulator (*CFTR*) chloride channel was strongly upregulated in PT-like CECs (Fig. 4G), whereas CD-like CECs showed minimal *CFTR* expression, with mean levels approximately sixfold lower than in PT-like CECs. The expression of *AVPR2*, which is an upstream activator of cAMP and encodes the direct molecular target of tolvaptan, was restricted to CD-like CECs (Fig. 4G). Immunofluorescence confirmed AVPR2 protein selectively in CD-like CECs and AQP2/LRP2 double-positive cells, which were not represented in our snRNA-seq dataset (Fig. 4H).

In summary, PT-like and CD-like CECs display distinct expression of molecular targets, suggesting differential amenability to interventions modulating redox balance, cAMP regulation, and solute-water transport. *CFTR* is enriched in PT-like CECs, whereas AVPR2 is restricted to CD-like CECs. Thus, the therapeutic impact of AVPR2 antagonism (tolvaptan) may depend on the burden of CD-like CECs within a patient’s cysts.

### Individual cyst type composition varies between patients

Because differential activation of molecular pathways in CECs may imply distinct therapeutic responsiveness, we assessed interpatient variability in cyst-type composition. We performed multicolor immunofluorescence for LRP2 and AQP2 on cystic tissue from 11 additional ADPKD patients (P1–P11) and analyzed 1,815 cysts to quantify PT-like, CD-like, and mixed cysts (Fig. 5A). To detect low-intensity marker expression without forcing assignments, we developed a machine-learning pipeline using the ilastik image-segmentation toolkit (*36*). Classifiers were trained via interactive annotation to recognize bright and dim epithelial fluorescence in three representative images per channel and then applied to all images. This approach unmasked LRP2 and AQP2 expression in many cysts with severely flattened epithelia (Suppl. Fig. 12). Final labels – PT-like, CD-like, mixed, or unclassified (LRP2^−^/AQP2^−^) – were assigned using both the model output and the original images (Suppl. Fig. 13).

**Figure 5.**
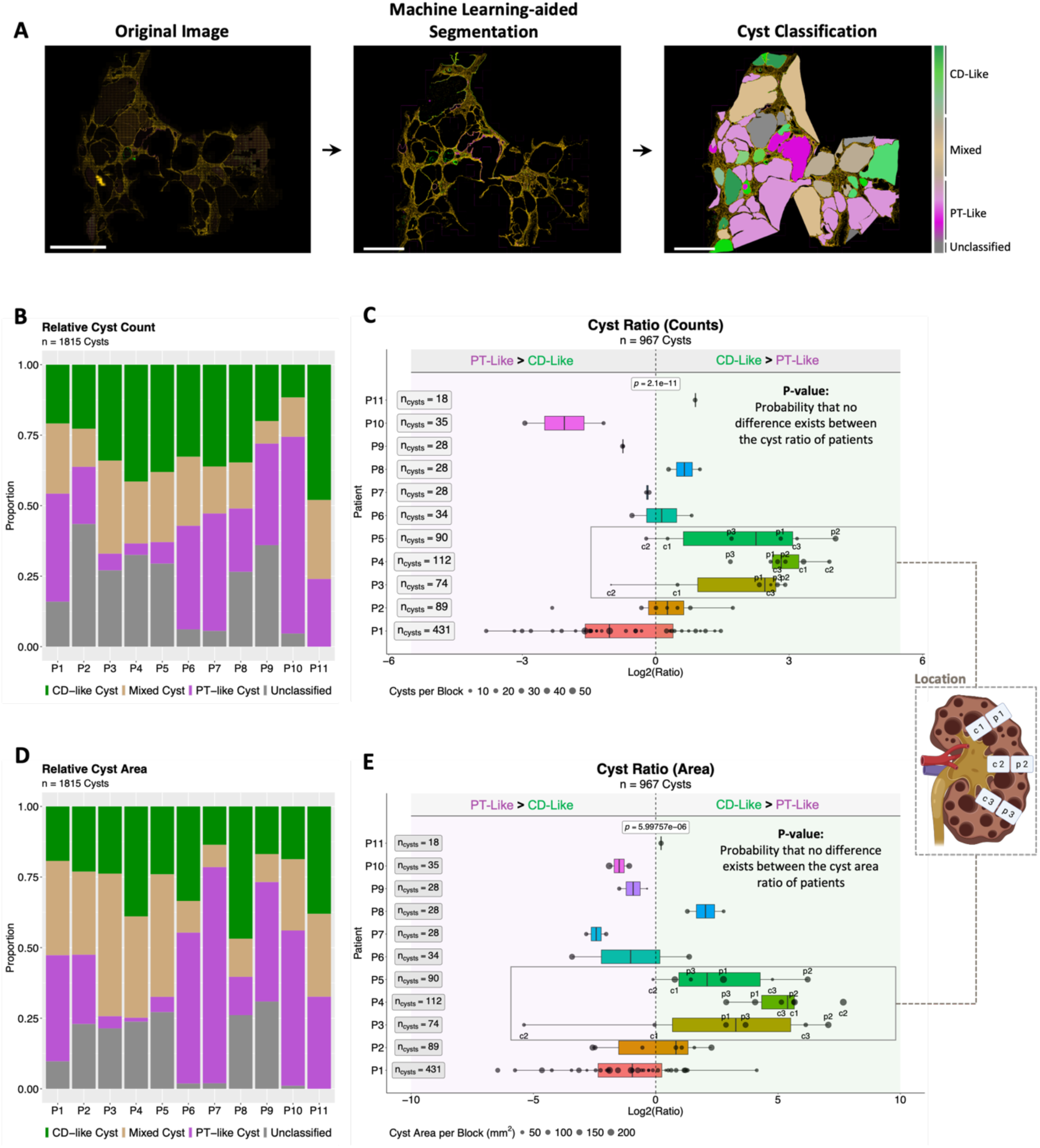
Individual cyst-type composition varies among ADPKD patients. (**A**) Analysis pipeline for cystic tissue using machine learning-aided image segmentation with ilastik. Scale bars, 5 mm. (**B**) Relative representation of cyst types per patient by cyst count. (**C**) Box plot of the log2-transformed ratio of CD-like to PT-like cysts within each analyzed kidney sample (paraffin section) by cyst count. (**D**) Relative representation of cyst types per patient by cumulative cyst area. (**E**) Box plot of the log2-transformed ratio of CD-like to PT-like cysts within each analyzed kidney sample by cyst area. Each dot represents one sample. A log2(ratio) < 0 indicates a bias toward PT-like cysts; a log2(ratio) > 1 indicates a bias toward CD-like cysts. Dot size reflects sample weight, i.e., the fraction of a patient’s total cyst count or area represented by the sample. Box plots show medians (center lines), 25th/75th percentiles (hinges), and whiskers extending 1.5× the interquartile range. nCysts indicates the sum of CD-like and PT-like cysts per patient. For patients P3–P5, sampling location is indicated (peripheral, p1–p3; central, c1–c3). P-values were calculated using the likelihood-ratio tests between linear mixed-effects models (Methods). Mixed and unclassified cysts were not included in CD-like/PT-like ratio analyses.

Across patients, cyst-type composition varied substantially: some kidneys were dominated by CD-like cysts, whereas others were enriched for PT-like cysts (Fig. 5B–E; Suppl. Fig. 15). The ratio of CD-like to PT-like cysts ranged from 1:3.5 (P10) to 8:1 (P4), consistent with prior puncture studies reporting variability in low-versus high-sodium cyst fluids across kidneys (*37*).

To evaluate potential spatial bias within large nephrectomy specimens, we generated whole-kidney center slices from patients P3–P5 and sampled spatially defined tissue blocks. We did not observe systematic differences in cyst-type composition between peripheral regions (near the renal capsule) and central regions (near the renal pelvis) (Suppl. Fig. 16).

Mixed cysts were detected in all patients (11/11) and accounted for 23% of total cyst number and 36% of total cyst area, indicating that they are a common feature of human ADPKD (Suppl. Fig. 17). On average, mixed cysts were also larger than PT-like or CD-like cysts (Suppl. Fig. 18). Overall, 24% of cysts were LRP2^−^/AQP2^−^ and therefore unclassified.

### Tolvaptan response is correlated with AQP2^+^ cyst burden in *Pkd1*^RC/RC^ mice

Because AVPR2 is exclusively expressed in AQP2^+^ CECs at both RNA and protein levels (Fig. 4G–H; Suppl. Fig. 14), differences in cyst-type composition may influence response to AVPR2 antagonism. However, tolvaptan has only been used in recent years so that long-term follow-up data and kidney nephrectomy samples are very scarce. Moreover, stratifying tolvaptan response in real-world cohorts is challenging due to a multitude of confounders, including comorbidities, therapy adherence, and lifestyle factors. We therefore leveraged a prior intervention study in which *Pkd1*^RC/RC^ mice received tolvaptan for 5 months and exhibited variable treatment responses by kidney weight to body weight ratio, cystic index, and cystic volume (*38*). Kidney sections from the three top responders and three top nonresponders (based on cystic volume) were stained for LRP2 and AQP2. Blinded quantification using our machine-learning-aided pipeline yielded per-animal cystic indices and cyst-type composition (Fig. 6A). All cyst types observed in human tissue – including LRP2^+^/AQP2^+^ mixed cysts – were present in *Pkd1*^RC/RC^ mice and the cystic index reliably recapitulated the response status of each animal (34) (Fig. 6B). Cyst classification showed that the abundance of CD-like cysts with high AQP2 expression (AQP2-high) was significantly associated with tolvaptan response, reflected by lower cystic indices. Because mixed cysts also contain AQP2^+^ CD-like CECs expressing AVPR2 (Fig. 4H; Suppl. Fig. 14), we next considered the total burden of AQP2^+^ cysts (CD-like plus mixed). Again, AQP2^+^ cyst abundance significantly correlated with treatment response. Taken together, these data support a model in which individual cyst type compositions modulate the responsiveness to tolvaptan.

**Figure 6.**
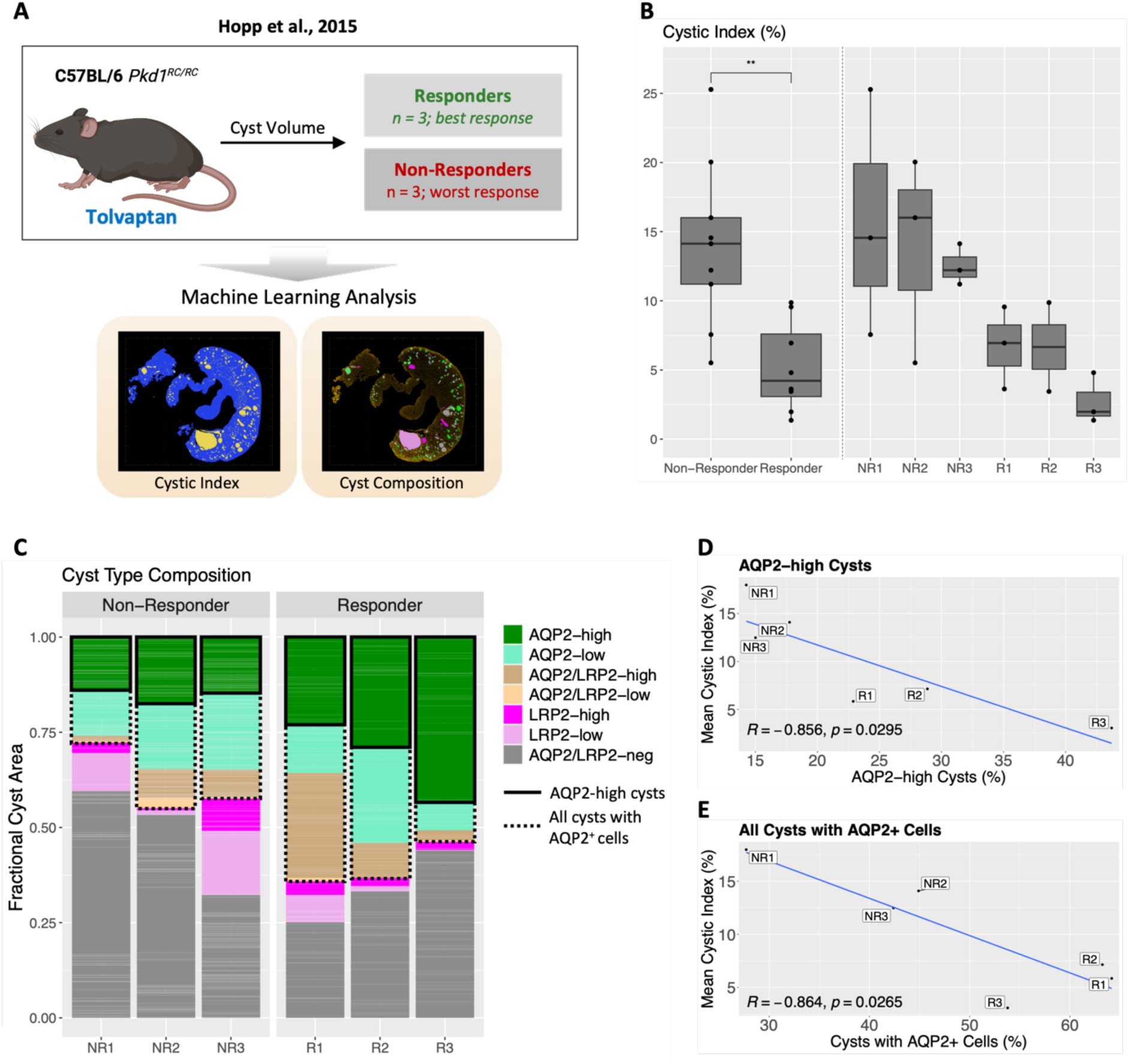
Abundance of AQP2^+^ cysts correlates with tolvaptan response in *Pkd1*^RC/RC^ mice. (**A**) Analysis pipeline for *Pkd1*^RC/RC^ kidney samples using machine-learning (ML)-aided image analysis. Kidney sections are from Hopp et al. (*38*). (**B**) Cystic index in tolvaptan responders and nonresponders based on ML-aided image analysis. (**C**) Relative representation of cysts expressing AQP2 (green/turquoise), LRP2 (magenta/pink), both markers (beige/brown), or neither marker (gray) in mice with good versus poor tolvaptan response. *, P ≤ 0.05; **, P ≤ 0.01; ***, P ≤ 0.001; ****, P ≤ 0.0001; no asterisk, P > 0.05.

## Discussion

We combined snRNA-seq with machine learning-guided image analysis of human ADPKD kidneys to resolve molecular cyst-type heterogeneity and its translational implications. Most cysts segregated into PT-like and CD-like classes based on epithelial lineage and cyst fluid composition. We further identified mixed cysts harboring LRP2^+^ and AQP2^+^ CECs as a common feature in ADPKD kidneys (Figs. 1, 5). These findings extend prior observations from a single-patient study that cells from adjacent nephron segments can co-occur within a cyst wall (*39*), showing that in mixed cysts, epithelial populations representing opposite ends of the nephron lie side-by-side as patches, mosaics, or even single interposed cells (Suppl. Fig. 5). This challenges the prevailing view that proximal and collecting duct markers are mutually exclusive in ADPKD cysts (*40*). Single nucleus transcriptomes stratified by epithelial lineage and cyst type further revealed that injury programs and glycolytic gene signatures are specifically amplified in mixed cyst epithelia. The origin of mixed cysts remains unresolved. Fusion of neighboring cysts seems unlikely, as we occasionally detected mononucleated double-positive cells during imaging (Fig. 4H, Suppl. Fig. 5). One possibility is aberrant LRP2 expression in subsets of differentiated collecting duct cells, for example due to somatic mutations. However, PT-like and CD-like CECs from mixed cysts clustered near control PT and CD principal cells, respectively, indicating broad transcriptomic similarity beyond individual markers (Suppl. Fig. 19). We therefore favor a model in which neighboring cells within a single tubular segment diverge along distinct differentiation trajectories. Consistent with this, dual LRP2/AQP2 staining was sometimes detectable in only mildly dilated tubules. Mixed cysts have not been widely reported in *in vivo* models, which remain the gold standard for investigating disease mechanisms (*41*). In the *Pkd1*^RC/RC^ mouse, we observed LRP2^+^/AQP2^+^ mixed cysts but at lower frequency than in human tissue. Systematic evaluation of mixed cyst prevalence across models and dissection of the mechanisms that generate them will be important for better understanding ADPKD pathophysiology.

Pathway analyses highlighted differential molecular targetability across cyst types (Fig. 4). Mixed cysts reside in a distinct microenvironment enriched for *CCL2* signaling, consistent with macrophage-mediated epithelial injury and accelerated cyst growth in ADPKD models, and with reports that CCL2 inhibition reduces inflammation and renal dysfunction in vivo (*25, 28, 42*). These data nominate CCL2 blockade as a strategy to curb growth of mixed cysts. Oxidative stress signatures were most pronounced in CD-like CECs, aligning with ROS accumulation in collecting ducts of *Pkd1*^−/−^ mice and with partial rescue via NRF2 activation (*31*). Restoring redox balance may therefore preferentially benefit CD-like cysts, a consideration for interpreting the negative phase 3 FALCON trial of the NRF2 activator bardoxolone (NCT03918447).

Prior work indicates that energy metabolism is rewired in cyst epithelia, with a shift toward glycolysis (*16*). Our snRNA-seq–based metabolic modeling suggests that this shift occurs predominantly in PT-like CECs, whereas energy demand and mitochondrial dysfunction are most pronounced in CD-like CECs (Fig. 2I-L). Together with repeated preclinical support for glycolysis inhibition (e.g., 2-deoxyglucose) and ketosis in ADPKD (*43–46*), these data highlight the need for cyst type–stratified evaluation of metabolic interventions, including dietary strategies.

Dysregulated cAMP signaling promotes cyst expansion (*3*). We delineate cyst type–specific differences in the expression of adenylyl cyclases, phosphodiesterases, and cAMP efflux transporters, and show unequal *CFTR* expression between PT-like and CD-like CECs. This heterogeneity may reconcile conflicting results on CFTR as a therapeutic target (46–48) and has implications for ongoing trials of CFTR inhibition (e.g., GLPG2737; NCT04578548). To date, AVPR2 inhibition remains the only approved therapy for ADPKD. Expression of AVPR2 was highly selective for CD-like CECs at both mRNA and protein levels, indicating that AVPR2-targeted agents act primarily on CD-like cysts, and partially on mixed cysts. In *Pkd1*^RC/RC^ mice, the magnitude of tolvaptan treatment response tracked with the abundance of AQP2^+^ cysts, supporting a link between cyst-type composition and AVPR2 pathway responsiveness and consistent with a mechanistic basis for interpatient variability in tolvaptan response.

The high burden of PT-like cysts we observed in some patients contrasts with the prevalent view that most cysts arise from the collecting duct. However, the predominance of high-sodium cyst fluids in puncture studies (*13, 37*) is consistent with a substantial PT-like contribution. In *Pkd1*^RC/RC^ mice, LRP2^+^ cysts reportedly predominate in early disease but decline over time, whereas AQP2^+^ cysts remain constant and LRP2/AQP2-negative cysts increase (*10*). We interpret this as progressive dedifferentiation reducing LRP2 detectability. Our image-analysis pipeline improved detection of low-intensity markers, enabling confident classification of more than 75% of cysts, including those with low LRP2 expression. Although some low-LRP2 cysts may originate from transition zones (e.g., PT–Bowman’s capsule or PT–descending thin limb), these findings argue that PT-like cysts merit explicit consideration in drug development, which often relies on collecting duct–focused models (*47*).

In this cohort, approximately 25% of cysts were LRP2^−^/AQP2^−^ by staining and likely derive from other nephron segments (e.g., thick ascending limb), which were underrepresented in the snRNA-seq dataset. Moreover, because tissue was obtained at nephrectomy from late-stage kidneys, early disease trajectories cannot be assessed directly. Access to early-stage human ADPKD tissue is inherently limited as patients are not nephrectomized and renal biopsy is rarely indicated, making molecular analysis of early-stage disease largely infeasible. Nevertheless, the analyzed patients were rapid progressors who typically meet criteria for disease-modifying therapy (*48*), underscoring clinical relevance. Larger cohorts and expanded marker panels will help to further refine cyst-type maps in ADPKD.

Noninvasive approaches to estimate global cyst-type composition in patients would facilitate stratified therapy. Sodium MRI (^23^Na-MRI) could quantify the burden of low-sodium, CD-like cysts across kidneys and inform the risk–benefit evaluation for AVPR2 antagonists, which can cause liver toxicity and polyuria. Because MRI is already used to track total kidney volume, adding ^23^Na-MRI may be feasible within current workflows.

In summary, we uncover previously underappreciated heterogeneity in cyst epithelial lineage, microenvironment, and pathway activation in human ADPKD, with direct relevance to therapeutic targeting. Accounting for cyst-type composition – especially the prevalence of mixed and PT-like cysts – may be essential for prioritizing targets, selecting models, and designing biomarker-guided trials aimed at improving outcomes in this heterogeneous disease.

## Materials and Methods

### Ethics statement

The use of kidney biopsies and nephrectomies in this study has been approved by the ethics committee of Charité Universitätsmedizin Berlin under the registration number EA4/026/18. The use of archived cystic kidney sections in of the validation cohort has been approved by the ethics committee of Hannover Medical School under the registration number 10183_BO_K_2022. All patients provided informed consent after oral and written elucidation on all aspects of the scientific project and their legal rights, including the right to withdraw the consent at any time without negative consequences. Patients were not offered any reward for tissue donation.

### Study cohort and collection of cysts and control tissues

We recovered a total of 15 superficial cysts from five end-stage ADPKD patients (3 cysts per patient) and three control samples of tumor adjacent normal kidney tissue from three control patients. All five ADPKD patients were on renal replacement therapy and undergoing total nephrectomy to create abdominal space prior to kidney transplantation. Control patients received total nephrectomy as part of their oncologic treatment. Kidney tissue and cyst specimens were stored in pre-cooled RNAlater at 4 °C for 24 h and then transferred to − 80 °C for further storage and snRNA-seq. None of the ADPKD patients received specific ADPKD treatment (tolvaptan) at the time of sample collection. Three of the five patients from the sequencing cohort had a mutation in the *PKD1* gene. For the other two patients the primary mutation was not known. The independent validation cohort comprised 67 archived samples (paraffin sections) from 11 additional ADPKD patients that underwent nephrectomy between February 2022 and Juli 2023 at Hannover Medical School. For 3 patients (P3-P5) the exact location of sample origin was known. Samples were archived in the Institute for Pathology at Hannover Medical School. Clinical patient data are summarized in **Supplementary Table 1**.

### Single nucleus dissociation and library preparation

Kidney tissue and ADPKD cyst specimens were processed for single-nuclei RNA sequencing as previously described (*49, 50*). In brief, samples were minced in nuclear lysis buffer 1 (nuclear lysis buffer (Sigma) + Ribolock (1U/µl) + VRC (10 mM)) and subsequently homogenized with a dounce homogenizer using pestle A (Sigma D8938-1SET). Samples were then filtered (100 µm strainer), re-homogenized (douncer with pestle B), and re-filtered (35-µm strainer). After centrifuging (5 min, 500 g), the pellet was resuspended in nuclear lysis buffer 2 (nuclear lysis buffer + Ribolock (1U/µl)). An underlay of 10% sucrose and 1U/µl of Ribolock in lysis buffer was used to remove debris in a second round of centrifugation (5 min, 500 g). Subsequently. supernatant and debris were carefully removed and pelleted nuclei resuspended in PBS / 0.04% BSA + Ribolock (1U/µl). Following a final filtration step (20-µm strainer) nuclei were stained with DAPI. All Specimens were kept at 4 °C throughout the procedure. Single-nuclei sequencing was performed according to the 10 × genomics protocol for Chromium Next GEM Single Cell 3’ v3.1 chemistry with a target count of 10,000 nuclei. Single nucleus RNA libraries were sequenced on Illumina HiSeq 4000 sequencers (paired end). Expression matrices were created using CellRanger (version 3.0.2) with --expect-cells 10,000 against the human HG38 genome (GRCh38 3.0.0). Background RNA was removed using CellBender.

### Bioinformatic data analysis

All integration and analysis steps were performed using the Seurat package in R (*8*). Sequenced nuclei with < 500 detected genes were removed during initial filtering. Before integration, the dataset of each sample was analyzed individually, including dimensionality reduction, unsupervised clustering, and projection in UMAP space following the Seurat standard workflow (https://satijalab.org/seurat/). Emerging clusters were then annotated for cell types the using expression of canonical marker genes using the FindAllMarkers function in Seurat (min.pct = 0.1, logfc.threshold = 1.5).

Individual datasets were then combined into an integrated dataset using the IntegrateLayers() function in Seurat version 5 (https://satijalab.org/seurat/) with the implemented harmony algorithm for batch correction (*9*). Subsequently, dimensionality reduction and unsupervised clustering were applied to the combined dataset. Cells with equal cell type labels from the individual dataset analysis clustered together in the integrated dataset, confirming successful integration and batch correction. Three of the 15 cyst samples were included in the integrated UMAP but removed from further downstream analyses and subclustering due to low recovery of cyst-lining epithelial cells (< 100 cells) in these samples. Destroyed nuclei and doublets were removed during initial clustering and subclustering, where they clustered away from the major cell types and/or showed expression of canonical markers from more than one major kidney cell type.

### Gene set enrichment analysis

The MSigDB hallmark gene sets (category H, https://www.gsea-msigdb.org) were used to identify overrepresented pathways in cell types and cyst types. Expression scores for each gene set were calculated per cell using UCell (*30*). Gene set expression heatmaps were created using the pheatmap tool in R.

### Calculation of energetic bias scores

The energy energetic bias score psi (𝜳_***E***_) was calculated as follows:

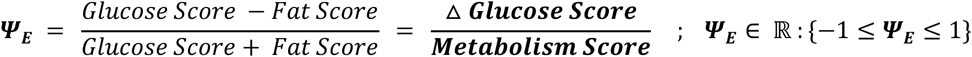

Genes summarized in the glucose score include core genes involved in glucose uptake (*SLC2A1*, *SLC2A2*, *SLC2A3*, *SLC2A4*), glycolysis (*HK1*, *PFKM*, *PKM*), and oxidative phosphorylation (*NDUFA6*, *SDHC*, *UQCRB*, *COX4I1*, *ATP5F1B*). Genes summarized in the fat score include core genes involved in fatty acid uptake (*CD36*, *FABP1*, *SLC27A2*), rate-limiting mitochondrial import (*CPT1A*, *CPT2*), β-oxidation (*ACADL*, *ACADM*, *ACADS*, *ECH1*, *ECHS1*, *EHHADH*, *HADH*, *HADHA*, *HADHB*, *ECI1*, and oxidative phosphorylation (*NDUFA6*, *SDHC*, *UQCRB*, *COX4I1*, *ATP5F1B*). The glucose and fat expression scores were calculated per cell using UCell in R (*30*). The total energy metabolism score was calculated as the sum of glucose and fat scores.

### Fluorescence microscopy and image data analysis of human samples

Samples were usually fixed in 4% formaldehyde at 4°C overnight and subsequently embedded in paraffin. Depending on the antibody used, formalin-fixed paraffin-embedded tissue slices were incubated with primary antibodies for 1h at room temperature or 4°C overnight. Incubation with secondary antibodies was performed at room temperature for 30min - 1h at room temperature. Cyst-lining epithelia of the sequencing cohort were stained using the anti-LRP2 monoclonal antibody [1:100, abcam, ab76969], the rabbit anti-AQP2 antibody [1:00; Thermo Fisher Scientifc, PA5-38004], or the goat anti-AQP2 antibody [1:100; Novus Biologicals, NBP1-70378]. For AVPR2 staining we used a rabbit anti-V2R antibody [1:100, Abcam, EPR24555-59]. For quantitative assessment of cyst type abundances in the validation cohort, we used the rabbit anti-aquaporin-2 [1:400, Sigma-Aldrich, 7310] and the mouse anti-LRP2 clone 10D5.1 [1:1000, Sigma-Aldrich, MABS489] as primary antibodies. As secondary antibodies we used donkey anti-mouse Alexa Fluor 555 [1:100, Invitrogen, A31570], donkey anti-rabbit Alexa Fluor 488 [1:100, Jackson Immunoresearch, 711-545-152], donkey anti-rabbit Alexa Fluor 488 [1:200, Invitrogen A21206], donkey anti-goat Alexa Fluor 488 [1:200, Life Technologies, A11055], and goat anti-rabbit Alexa Fluor 647 [1:200, Life Technologies, A21245]. Antibodies were diluted in PBS containing 5% BSA [Sigma A4503]. DNA was stained with mounting medium containing DAPI [Biozol, DNA-SCR-038448]. Slides were imaged using the Metafer Scanning Platform [MetaSystems Hard & Software, Altlussheim, Germany] at 40x or a Leica TCS SP8 confocal microscope equipped with a 63x PL APO CS 1.4 NA oil immersion objective lens [Leica, Wetzlar].

### Fluorescence microscopy and image data analysis of mouse samples

Formalin-fixed paraffin embedded kidney cross sections of C57BL/6J *Pkd1*^RC/RC^ mice treated with tolvaptan were chosen from the Hopp et al., J Am Soc Nephrol, 2015 study (*38*), based on treatment efficacy. Treatment efficacy was defined based on the reported cystic volume; the three animals with the highest (non-responders) and lowest (responders) cystic volume were evaluated independent of sex. For immunofluorescent staining kidneys underwent deparaffinization, antigen retrieval, and autofluorescence quenching as described previously (*51*). Proximal tubules/cysts were visualized with LRP2 (1:100, abcam, ab76969; 1:500, anti-mouse IgG1 Alexa Fluor-595, Invitrogen, A-21125) and collecting ducts/cyst were visualized with Aqp2 (1:100, Santa Cruz Biotechnology, sc-515770; 1:500, anti-rabbit IgG Alexa Fluor 488, Invitrogen, A-11034). Both, primary and secondary staining were performed for 1h at room temperature. All images were acquired using the Keyence BZ-X710.

### Cyst classification

Sequenced ADPKD cyst samples were classified as PT-like cysts or CD-like cysts, when snRNA-seq datasets contained only epithelial cells reflecting proximal or collecting duct segments, respectively. Cells were identified as proximal-like cyst epithelial cells (PT-like CECs) and collecting duct-like cyst epithelial cells (CD-like CECs) by expression of the PT marker LDL receptor-related protein 2 (*LRP2*) and the collecting duct marker aquaporin-2 (*AQP2*), respectively. Cysts that contained both *LRP2*^+^ and *AQP2*^+^ cells were classified as mixed cysts. Transcriptome-level cyst classification was confirmed using immunofluorescence staining for LRP2/megalin and AQP2 in formalin-fixed, paraffin-embedded sections of the sequenced cysts. Human samples of the validation cohort samples were classified based on immunofluorescence staining (LRP2, AQP2). Dilated tubule structures were defined as cysts based on the following criteria: *(i)* transverse diameter > 500µm and *(ii)* presence of epithelial cell lining. Dilated tubular structures with a pronouncedly mixed phenotype were also considered cysts. For mouse samples, were kidney samples and mean cyst size were considerably lower than in human ESRD kidneys, dilated tubule structures > 707µm^2^ (equivalent to a diameter > 30µm given a perfect circular shape) with an oblate ellipticity of > 0.15 were defined as cysts. Machine learning-base image segmentation samples was performed by training the image segmentation tool ilastik (*36*) through manual annotation of three different cystic tissue images per channel. The classifier was then applied to the rest of the images and signal probability map was exported. Because fully automated classification is sensitive to imaging artifacts, final cyst classes were assigned by evaluating, both, expression probability and original immunofluorescence signal. Cystic index calculations and cyst classification of mouse samples were performed blinded to disease severity and response to tolvaptan.

### Statistics

The significance of gene expression differences was evaluated using the Wilcoxon rank sum test within the stat_compare_means() function of the ggpubr package in R. The differences in cyst ratios between patients was evaluated by modelling log2 ratios with a linear mixed model using the lme4 package. The “patient” was used as a random factor to account for dependencies between each patient’s samples. Samples were weighted according to the proportion of a total patient’s cyst count or area represented in the respective sample. We compared the full model to a reduced model without patient as random factor: full.model = lmer(ratio.transformed ∼ 1 + (1 | Patient), weights = weights, data = data, REML = F). reduced.model = lm(ratio.transformed ∼ 1, weights = weights, data = data). The p-value was calculated using a Likelihood-Ratio-Tests: anova(reduced.model, full.model).

## List of Supplementary Materials

Suppl. Table 1

Suppl. Fig. 1 to Suppl. Fig. 19

## Acknowledgments

The authors thank Edda Christians, Tom Siol, and Anastasiya Boltengagen for excellent technical assistance. We would also like to acknowledge Cynthia Sieben, Peter Harris and the Mayo Clinic Robert M. and Billie Kelley Pirnie Translational Polycystic Kidney Disease (PKD) Center for providing the formalin-fixed paraffin-embedded kidney cross sections of C57BL/6J *Pkd1*^RC/RC^ mice treated with tolvaptan. Schematic illustrations were created with https://BioRender.com. Some of the dot plots were created with plot1cell (*52*).

## Authors contributions

JR performed bioinformatic data analysis, designed and performed analysis of microscopy data from sequencing and validation cohort, wrote the manuscript.

AG performed bioinformatic data analysis of sequencing cohort.

JK helped with the acquisition of cystic tissue samples from validation cohort

JS provided archived cystic tissue samples from validation cohort, performed staining and microscopic imaging.

FF recovered cysts from ADPKD kidney explants.

JB recovered cysts from ADPKD kidney explants.

RG helped with analysis of results.

MK helped in study design and analysis of results.

IAL performed staining of kidney sections from C57Bl6/J *Pkd1*^RC/RC^ mice

EWK helped in study design and analysis of results.

JH Performed genetic testing of patients from sequencing cohort and helped with analysis of results

KUE helped in study design.

CK help in designing the study and establishing the snRNA-seq for ADPKD samples.

JHB provided archived cystic tissue samples from validation cohort, performed staining and microscopic imaging.

RS helped in study design and analysis of results.

KH designed/executed staining experiments in C57Bl6/J *Pkd1*^RC/RC^ mice.

KSO helped in study design and analysis of results.

CH initiated, designed and supervised the study including bioinformatics analyses, performed initial sequencing experiments and wrote the manuscript.

## Data availability

Unfiltered Cellranger output files can be downloaded at NCBI GEO under the accession number GSEXXXXXX.

## References and Notes

1. M. B. Lanktree, A. Haghighi, E. Guiard, I.-A. Iliuta, X. Song, P. C. Harris, A. D. Paterson, Y. Pei, Prevalence Estimates of Polycystic Kidney and Liver Disease by Population Sequencing. Journal of the American Society of Nephrology : JASN 29, 2593–2600 (2018).

2. E. Cornec-Le Gall, A. Alam, R. D. Perrone, Autosomal dominant polycystic kidney disease. The Lancet 393, 919–935 (2019).

3. C. Bergmann, L. M. Guay-Woodford, P. C. Harris, S. Horie, D. J. M. Peters, V. E. Torres, Polycystic kidney disease. Nature Reviews Disease Primers 4 (2018), doi:10.1038/s41572-018-0047-y.

4. L. Luo, S. Roy, L. Li, M. Ma, Polycystic kidney disease: novel insights into polycystin function. Trends in molecular medicine 29, 268–281 (2023).

5. K. C. Yeung, E. Fryml, M. B. Lanktree, How Does ADPKD Severity Differ Between Family Members? Kidney International Reports 9, 1198–1209 (2024).

6. V. E. Torres, A. B. Chapman, O. Devuyst, R. T. Gansevoort, J. J. Grantham, E. Higashihara, R. D. Perrone, H. B. Krasa, J. Ouyang, F. S. Czerwiec, Tolvaptan in Patients with Autosomal Dominant Polycystic Kidney Disease. New England Journal of Medicine 367, 2407–2418 (2012).

7. I. Capuano, P. Buonanno, E. Riccio, M. Amicone, A. Pisani, Therapeutic advances in ADPKD: the future awaits. Journal of Nephrology 35, 397–415 (2022).

8. R. Satija, J. A. Farrell, D. Gennert, A. F. Schier, A. Regev, Spatial reconstruction of single-cell gene expression data. Nature Biotechnology 33, 495–502 (2015).

9. I. Korsunsky, N. Millard, J. Fan, K. Slowikowski, F. Zhang, K. Wei, Y. Baglaenko, M. Brenner, P. ru Loh, S. Raychaudhuri, Fast, sensitive and accurate integration of single-cell data with Harmony. Nature Methods 16, 1289–1296 (2019).

10. M. L. Nielsen, M. C. Mundt, D. L. Lildballe, M. Rasmussen, L. Sunde, V. E. Torres, P. C. Harris, H. Birn, Functional megalin is expressed in renal cysts in a mouse model of adult polycystic kidney disease. Clin. Kidney J. 14, 2420–2427 (2021).

11. X. Wang, L. Jiang, K. Thao, C. R. Sussman, T. LaBranche, M. Palmer, P. C. Harris, G. S. McKnight, K. P. Hoeflich, S. Schalm, V. E. Torres, Protein Kinase A Downregulation Delays the Development and Progression of Polycystic Kidney Disease. Journal of the American Society of Nephrology 33, 1087–1104 (2022).

12. K. Hopp, C. J. Hommerding, X. Wang, H. Ye, P. C. Harris, V. E. Torres, Tolvaptan plus pasireotide shows enhanced efficacy in a PKD1 model. Journal of the American Society of Nephrology 26, 39–47 (2015).

13. K. D. Gardner, Composition of Fluid in Twelve Cysts of a Polycystic Kidney. New England Journal of Medicine 281 (1969), doi:10.1056/NEJM1969103028118.

14. Y. Kirita, H. Wu, K. Uchimura, P. C. Wilson, B. D. Humphreys, Cell profiling of mouse acute kidney injury reveals conserved cellular responses to injury. Proceedings of the National Academy of Sciences of the United States of America 117, 15874–15883 (2020).

15. S. A. Yasinoglu, T. B. Kuipers, E. Suidgeest, L. van der Weerd, H. Mei, H. J. Baelde, D. J. M. Peters, Transcriptomic profiling of Polycystic Kidney Disease identifies paracrine factors in the early cyst microenvironment. Biochimica et Biophysica Acta - Molecular Basis of Disease 1870, 166987 (2024).

16. I. Rowe, M. Chiaravalli, V. Mannella, V. Ulisse, G. Quilici, M. Pema, X. W. Song, H. Xu, S. Mari, F. Qian, Y. Pei, G. Musco, A. Boletta, Defective glucose metabolism in polycystic kidney disease identifies a new therapeutic strategy. Nature Medicine 19, 488–493 (2013).

17. Y. Muto, E. E. Dixon, Y. Yoshimura, H. Wu, K. Omachi, N. Ledru, P. C. Wilson, A. J. King, N. Eric Olson, M. G. Gunawan, J. J. Kuo, J. H. Cox, J. H. Miner, S. L. Seliger, O. M. Woodward, P. A. Welling, T. J. Watnick, B. D. Humphreys, Defining cellular complexity in human autosomal dominant polycystic kidney disease by multimodal single cell analysis. Nature Communications 13, 1–19 (2022).

18. L. Wen, Y. Li, S. Li, X. Hu, Q. Wei, Z. Dong, Glucose Metabolism in Acute Kidney Injury and Kidney Repair. Frontiers in Medicine 8, 1–10 (2021).

19. W. Peng, C. Tan, L. Mo, J. Jiang, W. Zhou, J. Du, X. Zhou, X. Liu, L. Chen, Glucose transporter 3 in neuronal glucose metabolism: Health and diseases. Metabolism: clinical and experimental 123, 154869 (2021).

20. B. B. Lake, R. Menon, S. Winfree, Q. Hu, R. M. Ferreira, K. Kalhor, D. Barwinska, E. A. Otto, M. Ferkowicz, D. Diep, N. Plongthongkum, A. Knoten, S. Urata, A. S. Naik, S. Eddy, B. Zhang, Y. Wu, D. Salamon, J. C. Williams, X. Wang, K. S. Balderrama, P. Hoover, E. Murray, A. Vijayan, F. Chen, S. S. Waikar, S. Rosas, F. P. Wilson, P. M. Palevsky, K. Kiryluk, J. R. Sedor, R. D. Toto, C. Parikh, E. H. Kim, E. Z. Macosko, P. V. Kharchenko, J. P. Gaut, J. B. Hodgin, M. T. Eadon, P. C. Dagher, T. M. El-Achkar, K. Zhang, M. Kretzler, S. Jain, for the K. consortium, An atlas of healthy and injured cell states and niches in the human kidney. bioRxiv, 2021.07.28.454201 (2021).

21. J. J. Chung, L. Goldstein, Y. J. J. Chen, J. Lee, J. D. Webster, M. Roose-Girma, S. C. Paudyal, Z. Modrusan, A. Dey, A. S. Shaw, Single-cell transcriptome profiling of the kidney glomerulus identifies key cell types and reactions to injury. Journal of the American Society of Nephrology 31, 2341–2354 (2020).

22. B. R. Conway, E. D. O’Sullivan, C. Cairns, J. O’Sullivan, D. J. Simpson, A. Salzano, K. Connor, P. Ding, D. Humphries, K. Stewart, O. Teenan, R. Pius, N. C. Henderson, C. Bénézech, P. Ramachandran, D. Ferenbach, J. Hughes, T. Chandra, L. Denby, Kidney single-cell atlas reveals myeloid heterogeneity in progression and regression of kidney disease. Journal of the American Society of Nephrology 31, 2833–2854 (2020).

23. A. Puig-Kröger, E. Sierra-Filardi, A. Domínguez-Soto, R. Samaniego, M. T. Corcuera, F. Gómez-Aguado, M. Ratnam, P. Sánchez-Mateos, A. L. Corbí, Folate receptor β is expressed by tumor-associated macrophages and constitutes a marker for M2 anti-inflammatory/regulatory Macrophages. Cancer Research 69, 9395–9403 (2009).

24. S. Lee, S. Huen, H. Nishio, S. Nishio, H. K. Lee, B. S. Choi, C. Ruhrberg, L. G. Cantley, Distinct macrophage phenotypes contribute to kidney injury and repair. Journal of the American Society of Nephrology 22, 317–326 (2011).

25. M. F. Cassini, V. R. Kakade, E. Kurtz, P. Sulkowski, P. Glazer, R. Torres, S. Somlo, L. G. Cantley, Mcp1 promotes macrophage-dependent cyst expansion in autosomal dominant polycystic kidney disease. Journal of the American Society of Nephrology 29, 2471–2481 (2018).

26. T. Weimbs, Are cyst-associated macrophages in polycystic kidney disease the equivalent to tams in cancer? Journal of the American Society of Nephrology 29, 2447–2448 (2018).

27. S. Singh, D. Anshita, V. Ravichandiran, MCP-1: Function, regulation, and involvement in disease. International Immunopharmacology 101, 107598 (2021).

28. A. Viau, F. Bienaimé, K. Lukas, A. P. Todkar, M. Knoll, T. A. Yakulov, A. Hofherr, O. Kretz, M. Helmstädter, W. Reichardt, S. Braeg, T. Aschman, A. Merkle, D. Pfeifer, V. I. Dumit, M. Gubler, R. Nitschke, T. B. Huber, F. Terzi, J. Dengjel, F. Grahammer, M. Köttgen, H. Busch, M. Boerries, G. Walz, A. Triantafyllopoulou, E. W. Kuehn, Cilia-localized LKB 1 regulates chemokine signaling, macrophage recruitment, and tissue homeostasis in the kidney. The EMBO Journal 37, 1–21 (2018).

29. C. Kuppe, M. M. Ibrahim, J. Kranz, X. Zhang, S. Ziegler, J. Perales-Patón, J. Jansen, K. C. Reimer, J. R. Smith, R. Dobie, J. R. Wilson-Kanamori, M. Halder, Y. Xu, N. Kabgani, N. Kaesler, M. Klaus, L. Gernhold, V. G. Puelles, T. B. Huber, P. Boor, S. Menzel, R. M. Hoogenboezem, E. M. J. Bindels, J. Steffens, J. Floege, R. K. Schneider, J. Saez-Rodriguez, N. C. Henderson, R. Kramann, Decoding myofibroblast origins in human kidney fibrosis. Nature 589, 281–286 (2021).

30. M. Andreatta, S. J. Carmona, UCell: Robust and scalable single-cell gene signature scoring. Computational and Structural Biotechnology Journal 19, 3796–3798 (2021).

31. Y. Lu, Y. Sun, Z. Liu, Y. Lu, X. Zhu, B. Lan, Z. Mi, L. Dang, N. Li, W. Zhan, L. Tan, J. Pi, H. Xiong, L. Zhang, Y. Chen, Activation of NRF2 ameliorates oxidative stress and cystogenesis in autosomal dominant polycystic kidney disease. Science Translational Medicine 12, 1–16 (2020).

32. R. S. Balaban, S. Nemoto, T. Finkel, Mitochondria, oxidants, and aging. Cell 120, 483–495 (2005).

33. Y. R. Mehta, S. A. Lewis, K. T. Leo, L. Chen, E. Park, V. Raghuram, C. L. Chou, C. R. Yang, H. Kikuchi, S. Khundmiri, B. G. Poll, M. A. Knepper, “ADPKD-omics”: determinants of cyclic AMP levels in renal epithelial cells. Kidney International 101, 47–62 (2022).

34. K. P. Roos, K. A. Strait, K. L. Raphael, M. A. Blount, D. E. Kohan, Collecting duct-specific knockout of adenylyl cyclase type VI causes a urinary concentration defect in mice. American Journal of Physiology - Renal Physiology 302, 78–84 (2012).

35. F. Omar, J. E. Findlay, G. Carfray, R. W. Allcock, Z. Jiang, C. Moore, A. L. Muir, M. Lannoy, B. A. Fertig, D. Mai, J. P. Day, G. Bolger, G. S. Baillie, E. Schwiebert, E. Klussmann, N. J. Pyne, A. C. M. Ong, K. Bowers, J. M. Adam, D. R. Adams, M. D. Houslay, D. J. P. Henderson, Small-molecule allosteric activators of PDE4 long form cyclic AMP phosphodiesterases. Proceedings of the National Academy of Sciences of the United States of America 116, 13320–13329 (2019).

36. S. Berg, D. Kutra, T. Kroeger, C. N. Straehle, B. X. Kausler, C. Haubold, M. Schiegg, J. Ales, T. Beier, M. Rudy, K. Eren, J. I. Cervantes, B. Xu, F. Beuttenmueller, A. Wolny, C. Zhang, U. Koethe, F. A. Hamprecht, A. Kreshuk, Ilastik: Interactive Machine Learning for (Bio)Image Analysis. Nature Methods 16, 1226–1232 (2019).

37. R. Huseman, A. Grady, D. Welling, J. Grantham, Macropuncture study of polycystic disease in adult human kidneys. Kidney International 18, 375–385 (1980).

38. K. Hopp, C. J. Hommerding, X. Wang, H. Ye, P. C. Harris, V. E. Torres, Tolvaptan plus pasireotide shows enhanced efficacy in a PKD1 model. Journal of the American Society of Nephrology 26, 39–47 (2015).

39. Q. Li, Y. Wang, W. Deng, Y. Liu, J. Geng, Z. Yan, F. Li, B. Chen, Z. Li, R. Xia, W. Zeng, R. Liu, J. Xu, F. Xiong, C. L. Wu, Y. Miao, Heterogeneity of cell composition and origin identified by single-cell transcriptomics in renal cysts of patients with autosomal dominant polycystic kidney disease. Theranostics 11, 10064–10073 (2021).

40. O. Devuyst, C. R. Burrow, B. L. Smith, P. Agre, M. A. Knepper, P. D. Wilson, Expression of aquaporins-1 and -2 during nephrogenesis and in autosomal dominant polycystic kidney disease. American Journal of Physiology 271 (1996), doi:10.1152/ajprenal.1996.271.1.f169.

41. C. J. Sieben, P. C. Harris, Experimental Models of Polycystic Kidney Disease: Applications and Therapeutic Testing. Kidney360 4, 1155–1173 (2023).

42. C. Zoja, D. Corna, M. Locatelli, D. Rottoli, A. Pezzotta, M. Morigi, C. Zanchi, S. Buelli, A. Guglielmotti, N. Perico, A. Remuzzi, G. Remuzzi, Effects of MCP-1 inhibition by bindarit therapy in a rat model of polycystic kidney disease. Nephron 129, 52–61 (2015).

43. M. Chiaravalli, I. Rowe, V. Mannella, G. Quilici, T. Canu, V. Bianchi, A. Gurgone, S. Antunes, P. D’Adamo, A. Esposito, G. Musco, A. Boletta, 2-Deoxy-D-glucose ameliorates PKD progression. Journal of the American Society of Nephrology 27, 1958–1969 (2016).

44. R. Magistroni, A. Boletta, Defective glycolysis and the use of 2-deoxy-d-glucose in polycystic kidney disease: from animal models to humans. Journal of Nephrology 30, 511–519 (2017).

45. F. T. Chebib, K. L. Nowak, M. B. Chonchol, K. Bing, A. Ghanem, F. F. Rahbari-Oskoui, N. K. Dahl, M. Mrug, Polycystic Kidney Disease Diet: What is Known and What is Safe. Clinical Journal of the American Society of Nephrology 19, 664–682 (2024).

46. J. A. Torres, S. L. Kruger, C. Broderick, T. Amarlkhagva, S. Agrawal, J. R. Dodam, M. Mrug, L. A. Lyons, T. Weimbs, Ketosis Ameliorates Renal Cyst Growth in Polycystic Kidney Disease. Cell Metabolism 30, 1007–1023.e5 (2019).

47. T. B. Malas, W. N. Leonhard, H. Bange, Z. Granchi, K. M. Hettne, G. J. P. Van Westen, L. S. Price, P. A. C. ’t Hoen, D. J. M. Peters, Prioritization of novel ADPKD drug candidates from disease-stage specific gene expression profiles. EBioMedicine 51 (2020), doi:10.1016/j.ebiom.2019.11.046.

48. F. T. Chebib, R. D. Perrone, A. B. Chapman, N. K. Dahl, P. C. Harris, M. Mrug, R. A. Mustafa, A. Rastogi, T. Watnick, A. S. L. Yu, V. E. Torres, A practical guide for treatment of rapidly progressive ADPKD with tolvaptan. Journal of the American Society of Nephrology 29, 2458–2470 (2018).

49. J. Leiz, C. Hinze, A. Boltengagen, C. Braeuning, C. Kocks, N. Rajewsky, K. M. Schmidt-Ott, Nuclei isolation from adult mouse kidney for single-nucleus RNA-sequencing. Journal of Visualized Experiments 2021, 1–13 (2021).

50. C. Hinze, C. Kocks, J. Leiz, N. Karaiskos, A. Boltengagen, S. Cao, C. M. Skopnik, J. Klocke, J. H. Hardenberg, H. Stockmann, I. Gotthardt, B. Obermayer, L. Haghverdi, E. Wyler, M. Landthaler, S. Bachmann, A. C. Hocke, V. Corman, J. Busch, W. Schneider, N. Himmerkus, M. Bleich, K. U. Eckardt, P. Enghard, N. Rajewsky, K. M. Schmidt-Ott, Single-cell transcriptomics reveals common epithelial response patterns in human acute kidney injury. Genome Medicine 14, 1–18 (2022).

51. E. K. Kleczko, K. H. Marsh, L. C. Tyler, S. B. Furgeson, B. L. Bullock, C. J. Altmann, M. Miyazaki, B. Y. Gitomer, P. C. Harris, M. C. M. Weiser-evans, M. B. Chonchol, E. T. Clambey, R. A. Nemenoff, K. Hopp, CD8+ T cells modulate autosomal dominant polycystic kidney disease progression. Kidney International 94, 1127–1140.

52. H. Wu, R. G. Villalobos, X. Yao, D. Reilly, T. Chen, M. Rankin, E. Myshkin, M. D. Breyer, B. D. Humphreys, Mapping the single-cell transcriptomic response of murine diabetic kidney disease to therapies. Cell Metabolism 34, 1064–1078.e6 (2022).

