## Supplementary material for "Cyst-type epithelial heterogeneity shapes therapeutic responsiveness in ADPKD": Suppl. Figures

### Supplementary Figure Legends

Supplementary Table 1 | Patient characteristics within the sequencing and validation cohorts.

**Supplementary Figure 1 | Basic statistics of single-nucleus libraries. A**, Violin plots showing number of detected mRNA transcripts (nCount_RNA), genes (nFeature_RNA) as well as percentage of mitochondrial and ribosomal RNA per sample (percent.mt, percent.ribo). **B**, Bar plot depicting the absolute number of nuclei passing quality control. **C**, Relative representation of samples per cell type. **D**, Relative representation of patients per cell type. Adipo, adipocytes. CD-PC, collecting duct principal cells. CNT, connecting tubule. DCT, distal convoluted tubule. Endo, endothelial cells. Fibro, fibroblasts. IC-A, intercalated cells type A. IC-B, intercalated cells type B. Leuko, leukocytes. Lymph, lymphatic endothelial cells. PEC, parietal epithelial cells. Podo, podocytes. PT, proximal tubule. TAL, thick ascending limb. tL, thin limb.

**Supplementary Figure 2 | Quality control metrics and cell numbers in cell type clusters.** nCount_RNA, number of detected transcripts. nFeature_RNA, number of detected genes. percent.mt, percentage of mitochondrial transcripts. percent.ribo, percentage of ribosomal transcripts. Adipo, adipocytes. B, B cells. Baso/Mast, basophil granulocytes/mast cells. CD-PC, collecting duct principal cells. CD-like CECs, CD-like cyst epithelial cells. CNT, connecting tubule. DCT, distal convoluted tubule. Dmg, damaged cluster. DVR, descending vasa recta. EC, endothelial cell. Endo, endothelial cells. Fib, fibroblasts. Fibro, fibroblasts. Glom, glomerular endothelial cells. Healthy, healthy proximal tubule cells. IC-A, intercalated cells type A. IC-B, intercalated cells type B. Inflam, proinflammatory fibroblasts. Leuko, leukocytes. Lymph, lymphatic endothelial cells. Myo, myofibroblasts. PEC, parietal epithelial cells. Peri, pericytes. Plasma, plasma cells. Podo, podocytes. Prolif, proliferating cells. PT, proximal tubule. PT-like CECs, proximal tubule-leik cyst epithelial cells. T/NK, T cells/NK cells. TAL, thick ascending limb. tL, thin limb.

**Supplementary Figure 3 | Dimensionality reduction and canonical features. A**, UMAP reduction of integrated single nucleus RNA sequencing dataset of ADPKD and control samples. **B**, TSNE reduction of integrated single nucleus RNA sequencing dataset of ADPKD and control samples. **C**, Dot plot showing expression of canonical cell type markers in major cell type clusters. ADIPOQ, adiponectin. Adipo, adipocytes. AQP2, aquaporin-2. CALB1, calbindin 1. CD-PC, collecting duct principal cells. CDH5, cadherin 5. CIDEA, cell death-inducing DFFA-like effector A. CNT, connecting tubule. COL1A1, collagen type I alpha 1 chain. COL1A2, collagen type I alpha 2 chain. CUBN, cubilin. DCT, distal convoluted tubule. EPHA7, ephrin type-A receptor 7. Endo, endothelial cells. FCER1G, Fc fragment of IgE receptor Ig. Fibro, fibroblasts. FLT4, Fms related tyrosine kinase 4. FXYD4, FXYD domain-containing ion transport regulator 4. IC-A, intercalated cells type A. IC-B, intercalated cells type B. Leuko, leukocytes. LRP2, LDL receptor-related protein 2. Lymph, lymphatic endothelial cells. NPHS2, podocin. PEC, parietal epithelial cells. PECAM1, platelet endothelial cell adhesion molecule 1. PODXL, podocalyxin. Podo, podocytes. PROX1, prospero homeobox 1. PT, proximal tubule. PTPRC, protein tyrosine phosphatase, receptor type C. SLC12A1, solute carrier family 12 member 1. SLC12A3, solute carrier family 12 member 3. SLC26A4, solute carrier family 26 member 4. SLC26A7, solute carrier family 26 member 7. SLC4A1, solute carrier family 4 member 1. SLC4A9, solute carrier family 4 member 9. SLC44A5, solute carrier family 44 member 5. SLC8A1, solute carrier family 8 member 1. TAL, thick ascending limb. TRPM6, transient receptor potential cation channel subfamily M member 6. tL, thin limb. UMOD, uromodulin. VCAM1, vascular cell adhesion molecule 1.

**Supplementary Figure 4 | HE staining of cyst wall.** Formalin-fixed paraffin-embedded section of a cyst stained with hematoxilin-eosin (HE) showing general cyst wall architecture and distribution of epithelial cells, fibroblasts, immune cells, and endothelial cells. Scale bar, 100um.

**Supplementary Figure 5 | Distribution of PT-like CECs and CD-like CECs in mixed cysts.** Representative images showing the distribution of $LRP2^+$ PT-like CECs and $AQP2^+$ CD-like CECs in different mixed cysts. Arrowheads mark double positive cells. Scale bars, 100µm.

**Supplementary Figure 6 | Epithelial cells from mixed cysts display a pronounced injury phenotype. A**, Uniform manifold approximation and projection (UMAP) showing proximal epithelial cells coloured by their tissue of origin. **B**, Feature plots showing the expression of proximal tubule marker genes (upper row) as well as injury, chemotaxis, and repair genes (lower two rows). **C**, Heatmap illustrating the expression score of MSigDB hallmark gene sets. **D**, Uniform manifold approximation and projection (UMAP) showing collecting duct principal cells coloured by the tissue of origin. **E**, Feature plots showing the expression of collecting duct marker genes (upper row) as well as injury, chemotaxis, and repair genes (lower two rows). **F**, Heatmap illustrating the expression score of MSigDB hallmark gene sets. Gene sets were obtained from www.gsea-msigdb.org. Expression scores were calculated with UCell. Heatmaps were created using pheatmap. ACTA2, actin alpha 2. AQP2, aquaporin-2. AQP3, aquaporin-3. COL21A1, collagen type XXI alpha 1 chain. CXCL1, C-X-C motif chemokine ligand 1. CXCL2, C-X-C motif chemokine ligand 2. FBN, fibrillin. GADD45B, growth arrest and DNA damage inducible beta. HAVCR1, hepatitis A virus cellular receptor 1. LRP2, LDL receptor-related protein 2. NFKBIA, NFKB inhibitor alpha. SLC34A1, solute carrier family 34 member 1. SLC7A13, solute carrier family 7 member 13. TNFRSF12A, TNF receptor superfamily member 12A. VCAM1, vascular cell adhesion molecule 1.

**Supplementary Figure 7 | Epithelial cells from mixed cysts display a pronounced metabolic shift.** Dot plots and violin plots illustrating the expression of key genes involved in fatty acid uptake (upper panels) and beta oxidation (lower panels) per tissue type in proximal epithelial cells and collecting duct epithelial cells. ACADL, acyl-CoA dehydrogenase, long chain. ACADM, acyl-CoA dehydrogenase, medium chain. ACADS, acyl-CoA dehydrogenase, short chain. CPT1A, carnitine palmitoyltransferase 1A. CPT2, carnitine palmitoyltransferase 2. ECI1, enoyl-CoA delta isomerase 1. EHHADH, enoyl-CoA hydratase and 3-hydroxyacyl CoA dehydrogenase. FABP1, fatty acid binding protein 1. HADH, hydroxyacyl-CoA dehydrogenase. HADHA, hydroxyacyl-CoA dehydrogenase trifunctional multienzyme complex subunit alpha. HADHB, hydroxyacyl-CoA dehydrogenase trifunctional multienzyme complex subunit beta. SLC27A2, solute carrier family 27 member 2. *: $p \leq 0.05$ **: $p \leq 0.01$, ***: $p \leq 0.001$, ****: $p \leq 0.0001$. No asterisks: $p > 0.05$.

**Supplementary Figure 8 | PT-like CECs stop shuttling glucose and amino acids and shut down gluconeogenesis. A**, Dot plots and Violin plots showing the expression of key genes involved in proximal tubule glucose shuttling (**A**), gluconeogenesis (**B**), apical amino acid transport (**C**), and basolateral amino acid transport (**D**). Overlayed bar plots indicate mean expression. Genes were selected according to Makrides et al., Transport of Amino Acids in the Kidney, Compr. Physiol. (2014). FBP1, fructose-bisphosphatase 1. G6PC, glucose-6-phosphatase, catalytic subunit 1. PC, pyruvate carboxylase. PCK1, phosphoenolpyruvate carboxykinase 1. SLC15A1, solute carrier family 15 member 1. SLC15A2, solute carrier family 15 member 2. SLC16A10, solute carrier family 16 member 10. SLC1A1, solute carrier family 1 member 1. SLC2A1, solute carrier family 2 member 1. SLC2A2, solute carrier family 2 member 2. SLC3A1, solute carrier family 3 member 1. SLC3A2, solute carrier family 3 member 2. SLC36A1, solute carrier family 36 member 1. SLC36A2, solute carrier family 36 member 2. SLC43A2, solute carrier family 43 member 2. SLC5A1, solute carrier family 5 member 1. SLC5A2, solute carrier family 5 member 2. SLC6A18, solute carrier family 6 member 18. SLC6A19, solute carrier family 6 member 19. SLC6A20, solute carrier family 6 member 20. SLC7A7, solute carrier family 7 member 7. SLC7A8, solute carrier family 7 member 8. SLC7A9, solute carrier family 7 member 9. *: $p \leq 0.05$ **: $p \leq 0.01$, ***: $p \leq 0.001$, ****: $p \leq 0.0001$. No asterisks: $p > 0.05$.

**Supplementary Figure 9 | Marker gene expression in cell types of the cyst microenvironment. A-C**, Violin plots depicting the expression of marker genes in subclusters of immune cells, fibroblasts, and endothelial cells. Horizontal bars represent mean expression levels. ACTA2, actin alpha 2. AQP1, aquaporin 1. B, B cells. Baso/Mast, basophil granulocytes/mast. CD19, CD19 molecule. CD3E, CD3e molecule. CD4, CD4 molecule. CD8A, CD8a molecule. CD36, CD36 molecule. CDK1, cyclin-dependent kinase 1. COL1A1, collagen type I alpha 1 chain. COL1A2, collagen type I alpha 2 chain. COL4A3, collagen type IV alpha 3 chain. COL4A4, collagen type IV alpha 4 chain. Dmg, damaged cluster. DVR, descending vasa recta. EC, endothelial cells. EDH3, EH domain containing 3. FCER1A, Fc epsilon receptor Ia. Glom, glomerular endothelial cells. HECW2, HECT, C2 and WW domain containing E3 ubiquitin protein ligase 2. IGFBP3, insulin-like growth factor binding protein 3. IGFBP5, insulin-like growth factor binding protein 5. IGFBP7, insulin-like growth factor binding protein 7. IGHA1, immunoglobulin heavy constant alpha 1. IGHG1, immunoglobulin heavy constant gamma 1. IL3RA, interleukin 3 receptor subunit alpha. IL5RA, interleukin 5 receptor subunit alpha. IL6, interleukin 6. Inflam, proinflammatory fibroblasts. ITGAX, integrin subunit alpha X. MEG3, maternally expressed 3. MKI67, marker of proliferation Ki-67. MS4A1, membrane-spanning 4-domains A1. Myo, myofibroblasts. NKD2, NKD inhibitor of WNT signaling pathway 2. NOTCH3, notch receptor 3. NPR3, natriuretic peptide receptor 3. PDGFRA, platelet-derived growth factor receptor alpha. PDGFRB, platelet-derived growth factor receptor beta. PECAM1, platelet endothelial cell adhesion molecule 1. Plasma, plasma cells. PLVAP, plasmalemma vesicle associated protein. POSTN, periostin. Prolif, proliferating cells. PTPRC, protein tyrosine phosphatase receptor type C. SLC14A1, solute carrier family 14 member 1. T/NK, T cells/NK cells.

**Supplementary Figure 10 | Relative abundance of kidney cell types. A,** Fraction of major kidney cell types relative to the number of sequenced cells per tissue type. **B,** Fraction of major kidney cell types relative to the number of sequenced cells in control (C1-C3) and ADPKD (P1-P5) patients. Podo, podocytes. PEC, parietal epithelial cells. PT, proximal tubule. tL, thin limb. TAL, thick ascending limb. DCT, distal convoluted tubule. CNT, connecting tubule. CD-PC, collecting duct principal cells. IC-A, intercalated cells type A. IC-B, intercalated cells type B. Endo, endothelial cells. Lymph, lymphatic endothelial cells. Adipo, adipocytes. Leuko, leukocytes. Fibro, fibroblasts.

**Supplementary Figure 11 | CD-like CECs from mixed cysts carry the highest oxidative stress burden.** Overlay of violin pots and box plots showing the expression of the MSigDB hallmark reactive oxygen species pathway gene set per cell type and tissue type. Expression scores were calculated using UCell. No asterisks: $p > 0.05$, *: $p \leq 0.05$ **: $p \leq 0.01$, ***: $p \leq 0.001$, ****: $p \leq 0.0001$.

**Supplementary Figure 12 | Machine learning-aided imaging segmentation unmasks dimmed marker gene expression.** Representative images illustrating the microscopic marker gene detection in the original immunofluorescence image and detection in the ilastik segmented image for (**A**) bright fluorescence intensities and (**B**) low fluorescence intensities. Scale bars, 500µm

**Supplementary Figure 13 | Microscopic cyst classes.** Representative images showing LRP2 and AQP2 staining in cysts of different classes. +++ = strong expression, + = dimmed expression, (+) = expression cannot be excluded. Scale bars, 200µm.

**Supplementary Figure 14 | Selective protein expression of AVPR2 in cysts lined with AQP2$^{+}$ CD-like CECs. A**, Immunofluorescence image showing expression of AQP2, LRP2, and AVPR2 in biopsy tissue from kidney allografts. Nuclei are stained with DAPI. **B**, Cyst classes following machine learning-aided image segmentation. **C**, Immunofluorescence staining showing distribution of AVPR2 expression in the cyst epithelium of different cyst types. Filled arrowheads indicate AVPR2 epithelia, hollow arrowheads mark epithelia without AVPR2 expression. Scale bars, 1000µm (overview) and 100µm (inset).

**Supplementary Figure 15 | Detailed cyst classification.** Stacked bar plots showing relative representation of fine and coarse microscopic cyst classes per sample and patient, regarding cyst count (**A, C**) and cyst area (**B, D**). Each stacked bar plot represents one kidney sample. Diamond symbols indicate the weight of each sample, i.e., the fraction of a patient's total cyst count or cyst area represented within the respective sample.

**Supplementary Figure 16 | Cyst classes per location.** Stacked bar plots showing relative representation of fine and coarse microscopic cyst classes per sample and location (peripheral vs. central), regarding cyst count (**A, C**) and cyst area (**B, D**). Each stacked bar plot represents one kidney sample. Significance was tested using a Likelihood-Ratio-Test between a linear mixed-effect model with "location" as random factor and a reduced linear model without this random factor. n.s., not significant.

**Supplementary Figure 17 | Overall relative representation of cyst type classes.**

**Supplementary Figure 18 | Cyst size per cyst type.** Boxplots showing cyst area per cyst type. Central lines show median values. Lower and upper hinges indicate 25%-quartiles and 75%-quartiles, respectively. Upper and lower whiskers extend 1.5 * inter-quartile range from the respective hinge. *: $p \leq 0.05$ **: $p \leq 0.01$, ***: $p \leq 0.001$, ****: $p \leq 0.0001$. No asterisks: $p > 0.05$.

**Supplementary Figure 19 | Distribution of PT-like and CD-like CECs from mixed cysts on integrated UMAP.**

### Supplementary Table 1

**Sequencing Cohort**

| Patient | Group | Sex | Mutation | Age at Nephrectomy | GFR at Nephrectomy |
|---|---|---|---|---|---|
| C1 | Control | F | N/A | 51 | 55 |
| C2 | Control | M | N/A | 61 | 60 |
| C3 | Control | M | N/A | 53 | 73 |
| P1 | PKD | M | *PKD1*: c.1297C>T, p.(Gln433*), heterozygous | 46 | RRT |
| P2 | PKD | M | unknown | 45 | RRT |
| P3 | PKD | M | *PKD1: c.8299C>G, p.(Arg2767Gly), heterozygous* | 46 | RRT |
| P4 | PKD | F | *PKD1: c.5608A>C, p.(Asn1870His), heterozygous* | 56 | RRT |
| P5 | PKD | M | unknown | 59 | RRT |

**Validation Cohort**

| Patient | Group | Sex | Mutation | Age at Nephrectomy | GFR at Nephrectomy |
|---|---|---|---|---|---|
| P1 | PKD | F | unknown | 74 | RRT |
| P2 | PKD | M | unknown | 53 | RRT |
| P3 | PKD | F | unknown | 70 | RRT |
| P4 | PKD | F | unknown | 64 | RRT |
| P5 | PKD | M | unknown | 61 | RRT |
| P6 | PKD | F | unknown | 59 | RRT |
| P7 | PKD | M | unknown | 55 | RRT |
| P8 | PKD | M | unknown | 47 | RRT |
| P9 | PKD | F | unknown | 67 | RRT |
| P10 | PKD | M | unknown | 63 | RRT |
| P11 | PKD | M | unknown | 58 | RRT |

### Supplementary Figure 1

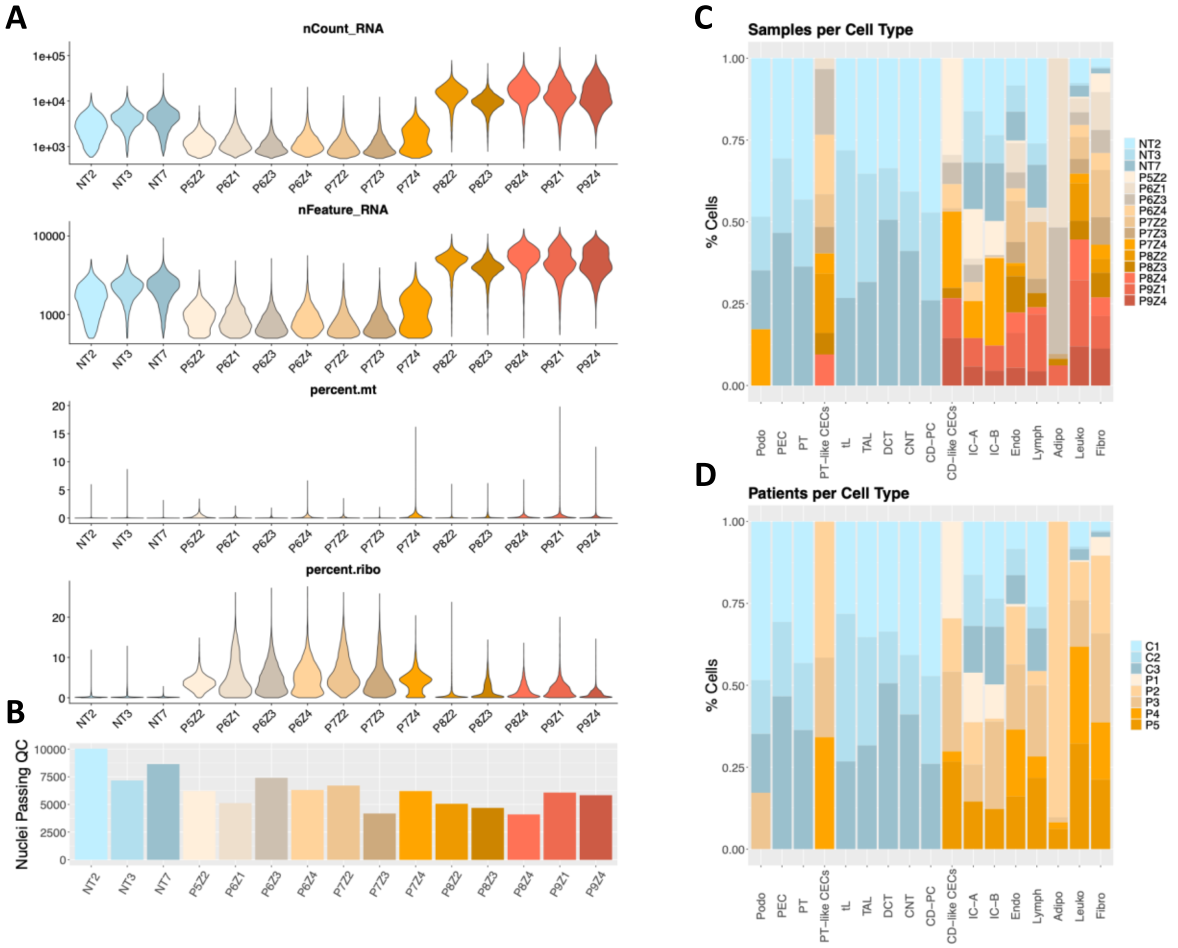

### Supplementary Figure 2

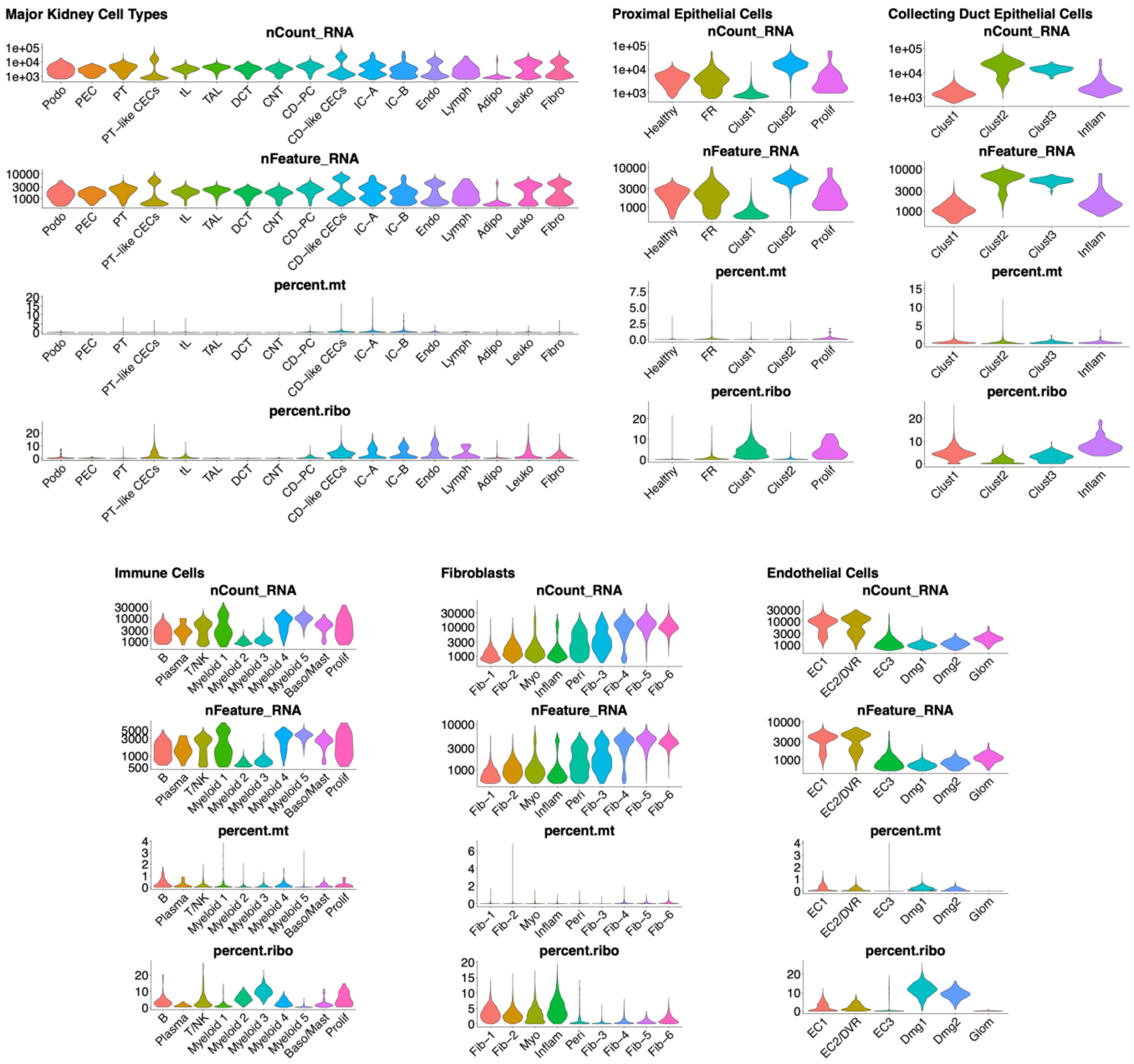

Supplementary Figure 3

A

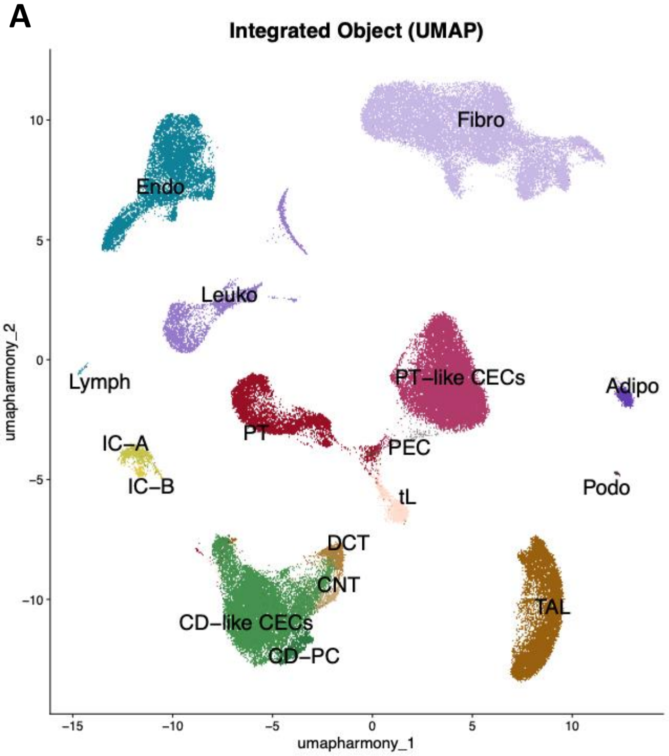


B

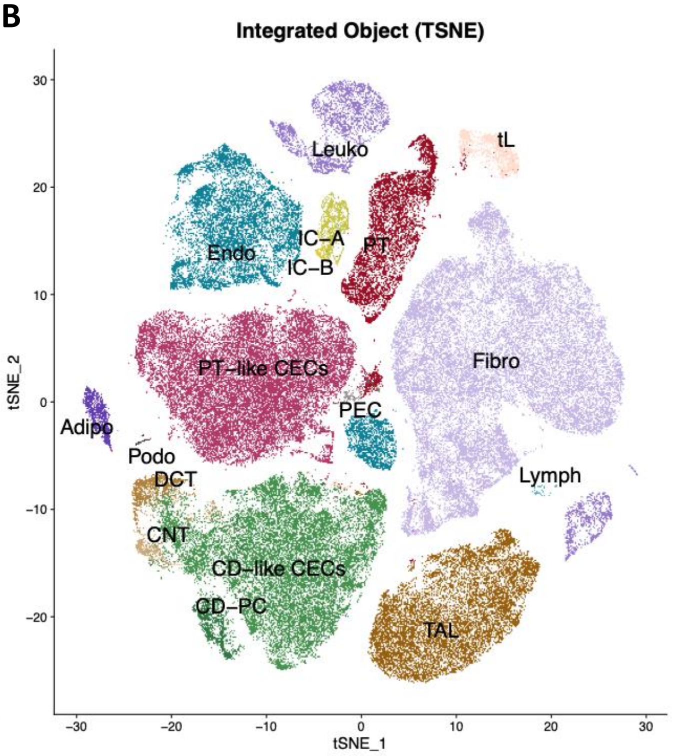


C

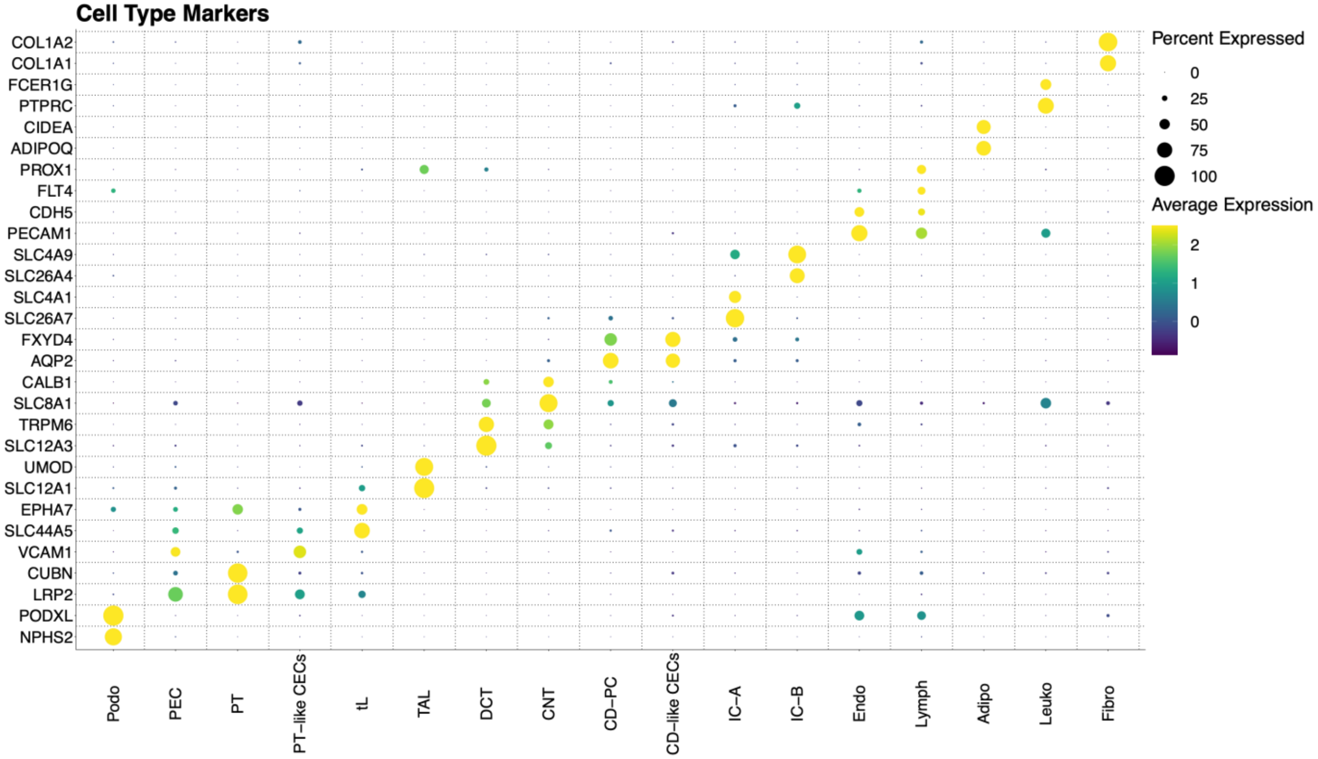

Supplementary Figure 4

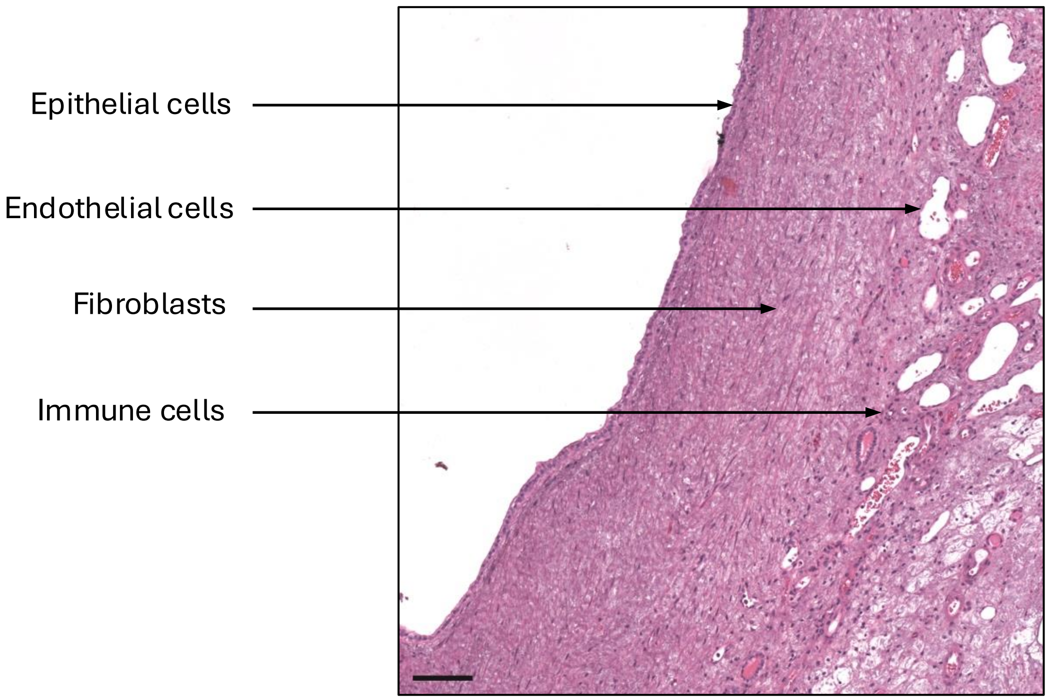

Supplementary Figure 5

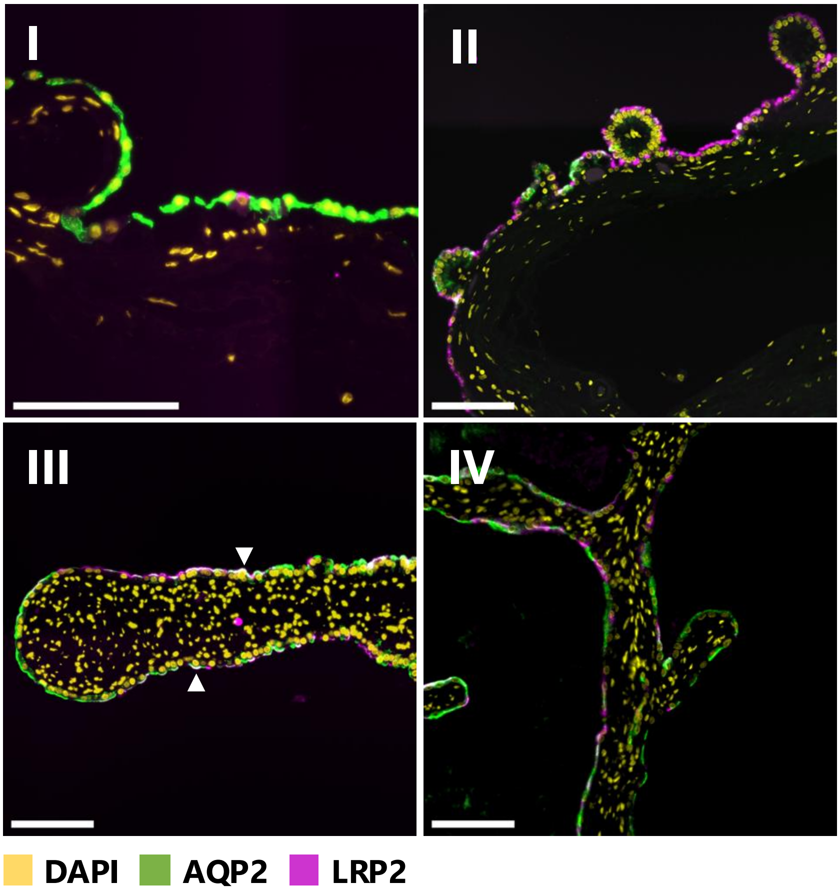

### Supplementary Figure 6

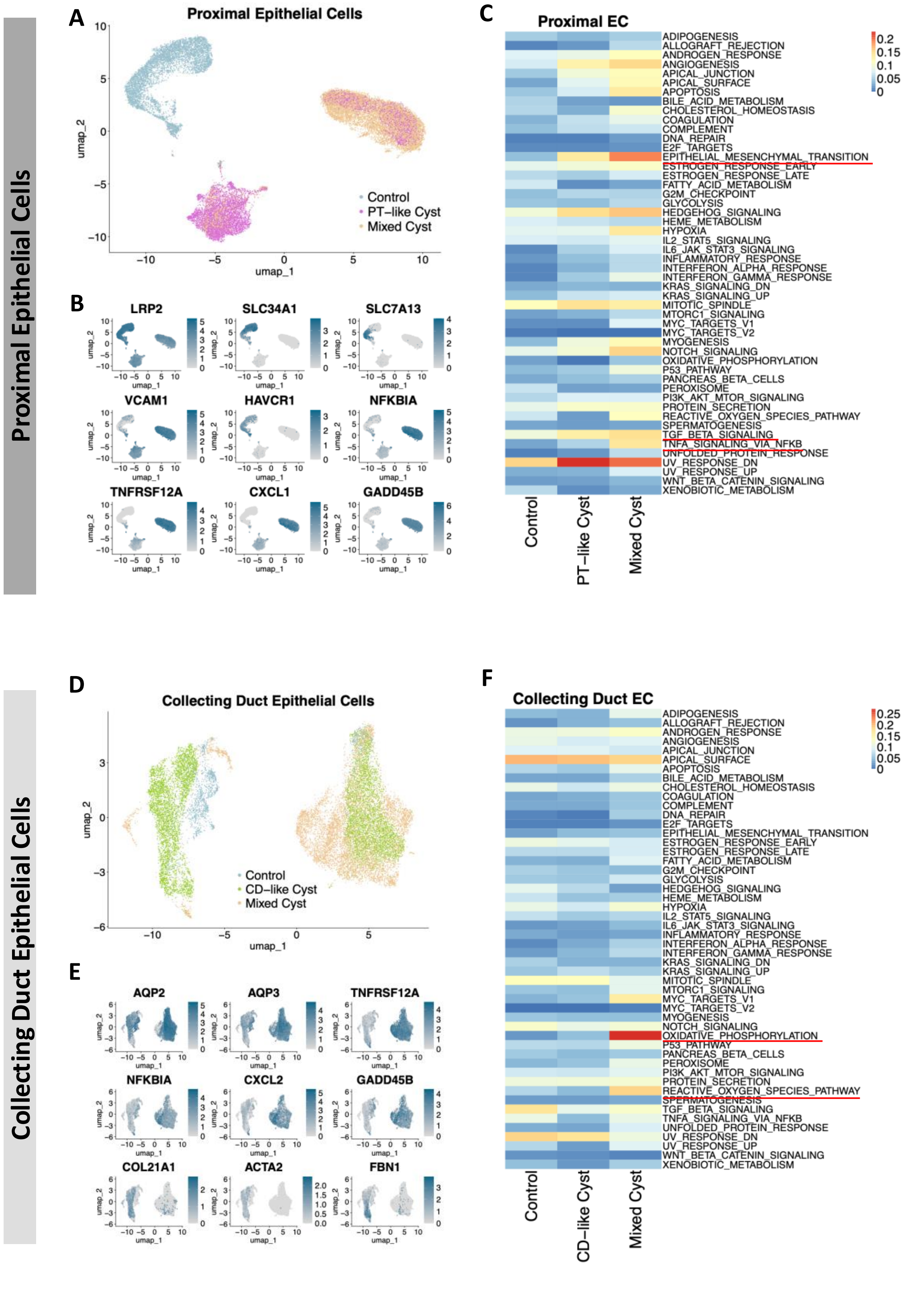

### Supplementary Figure 7

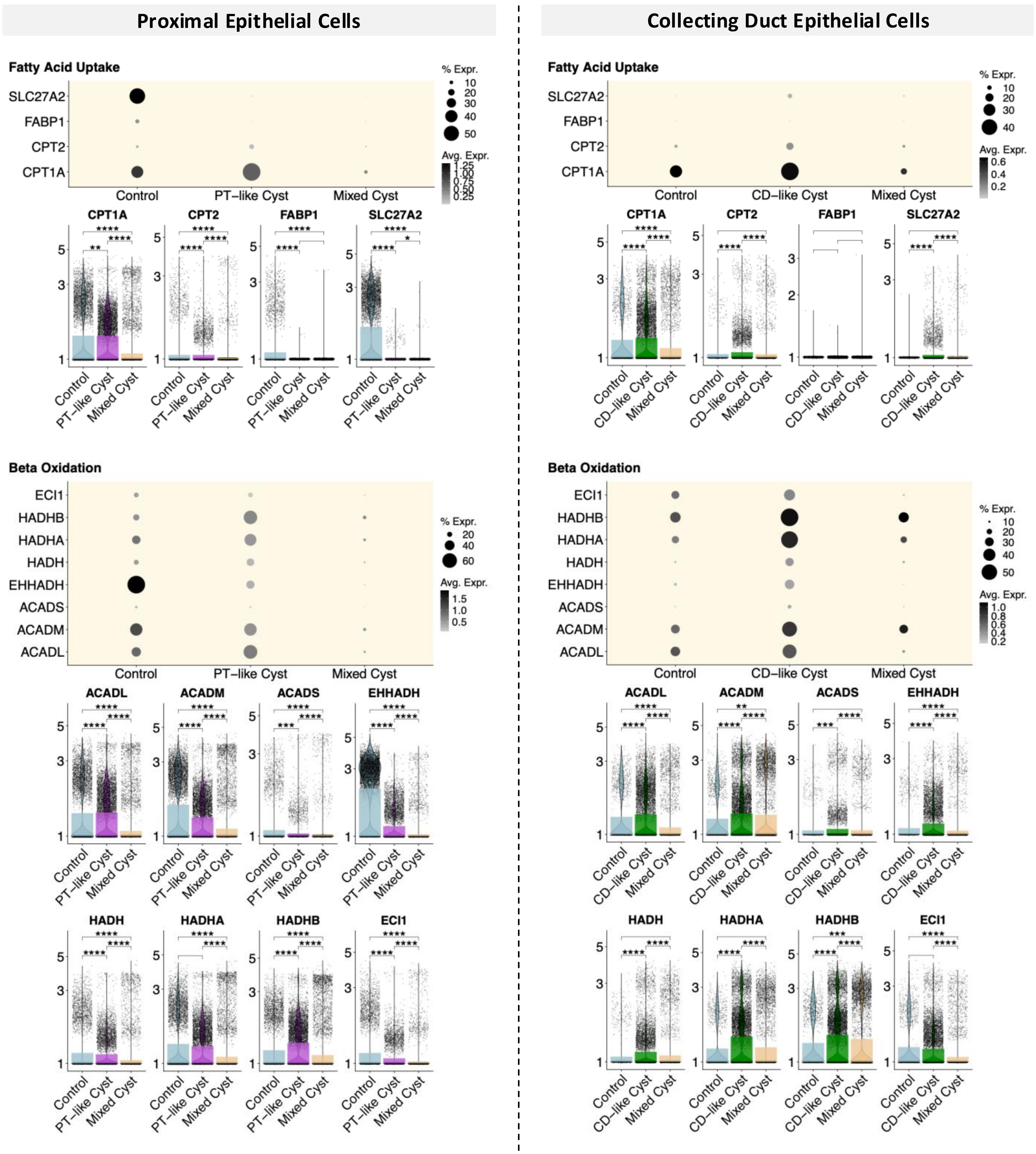

### Supplementary Figure 8

**A**

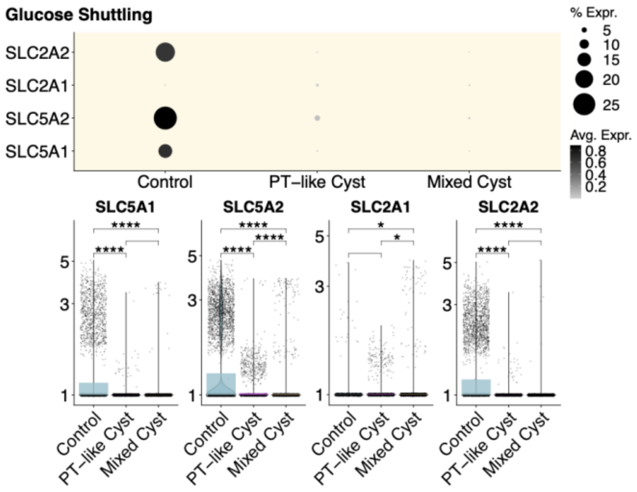


**B**

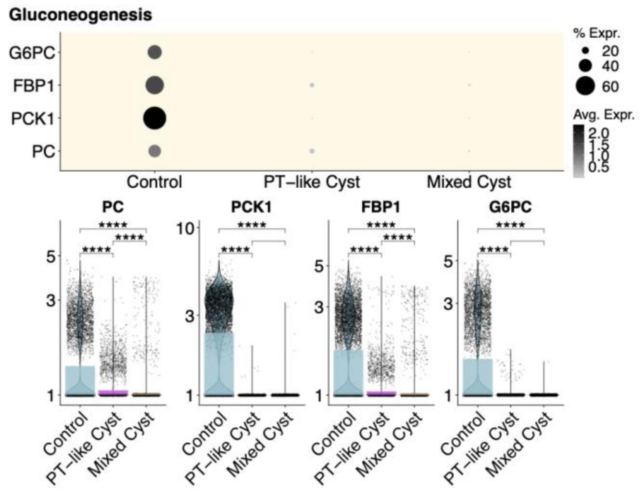


**C**

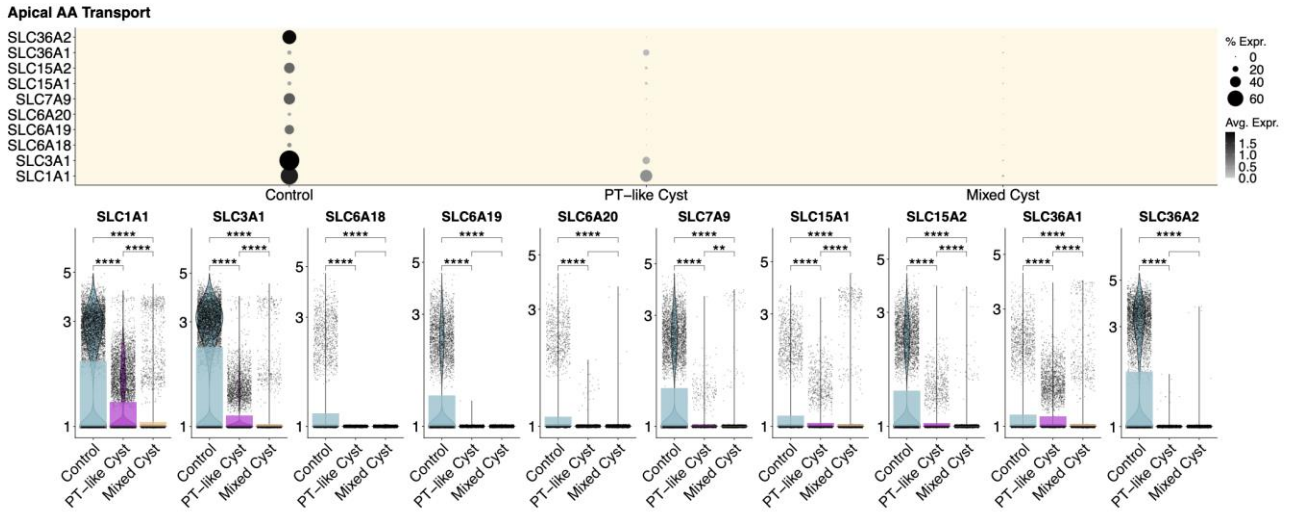


**D**

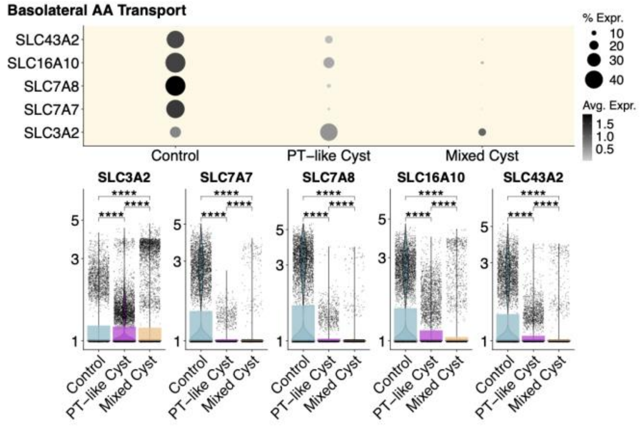

Supplementary Figure 9

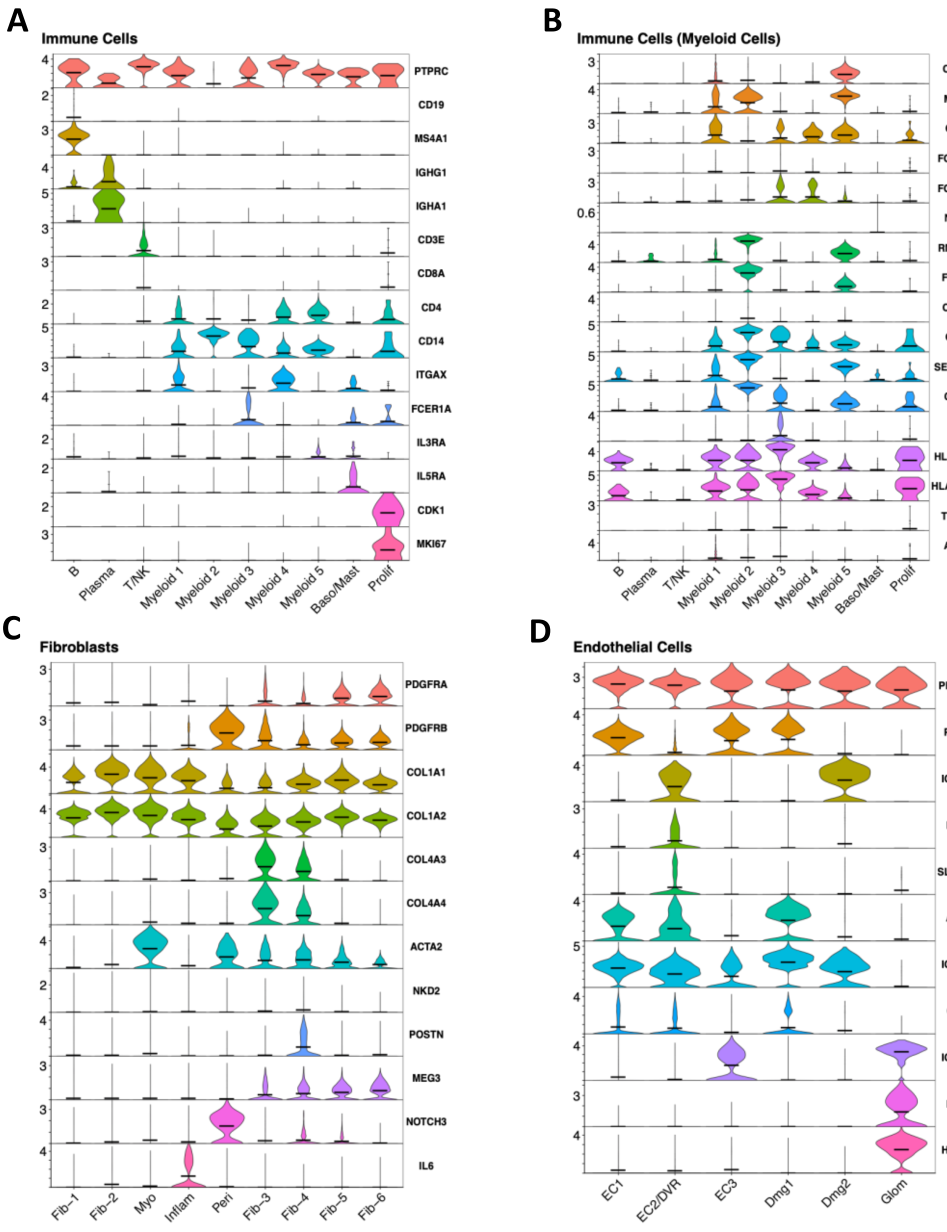

Supplementary Figure 10

**A**

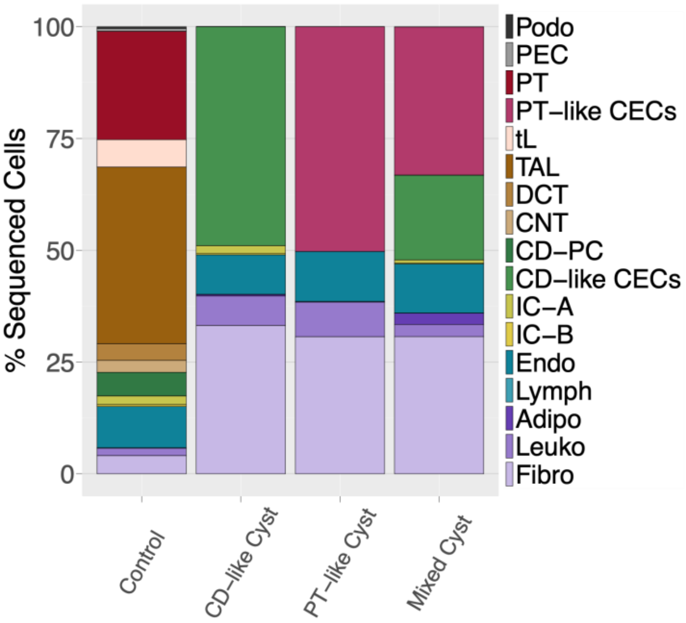


**B**

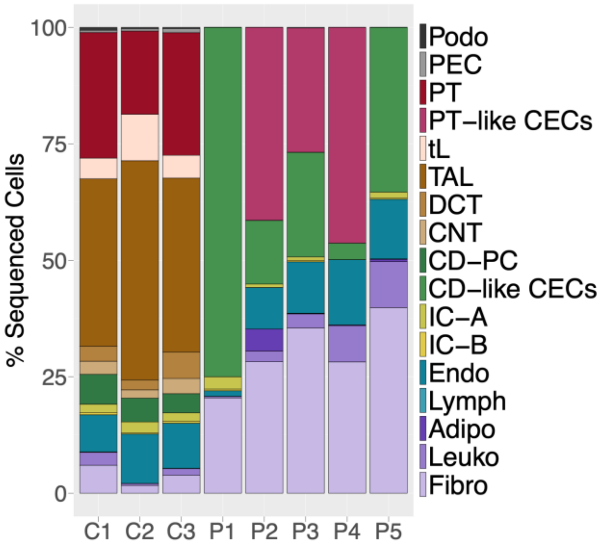

Supplementary Figure 11

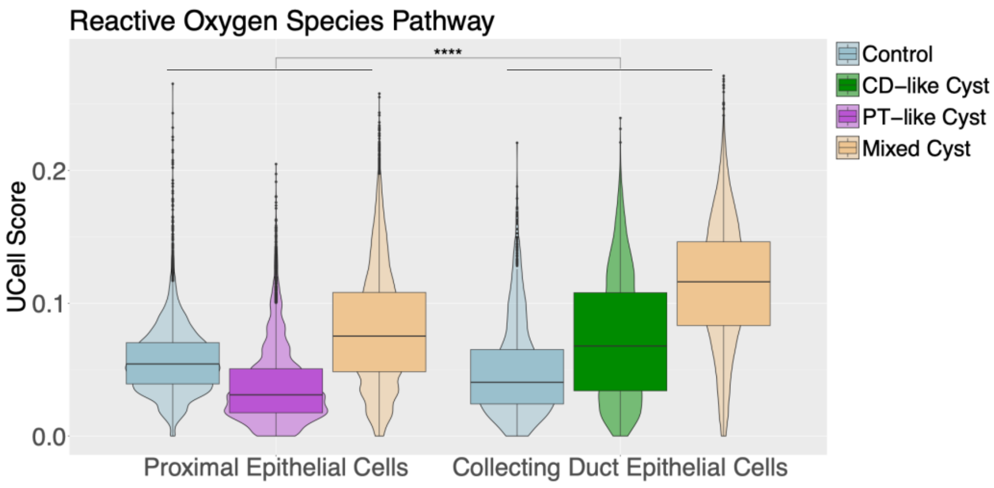

Supplementary Figure 12

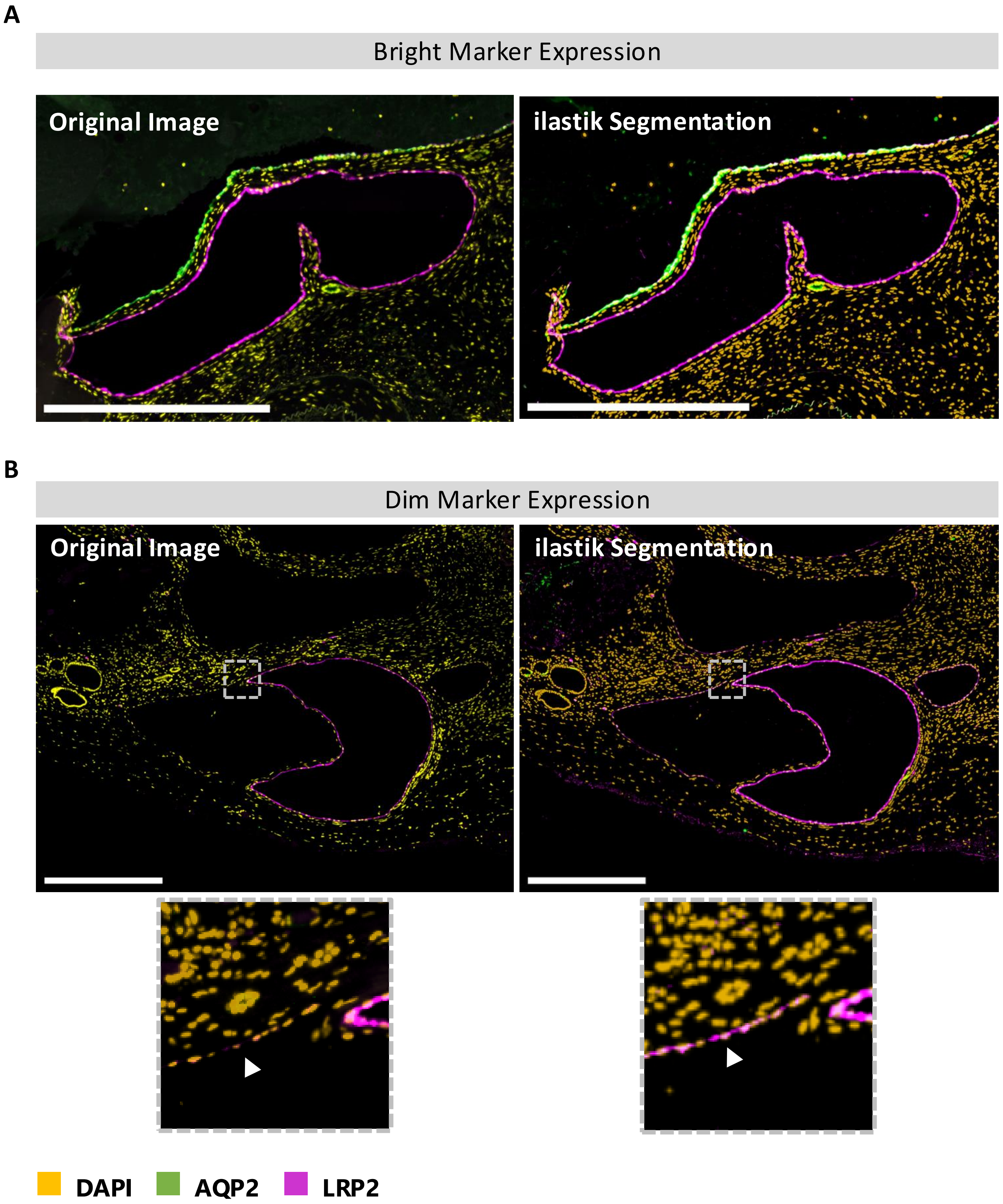

### Supplementary Figure 13

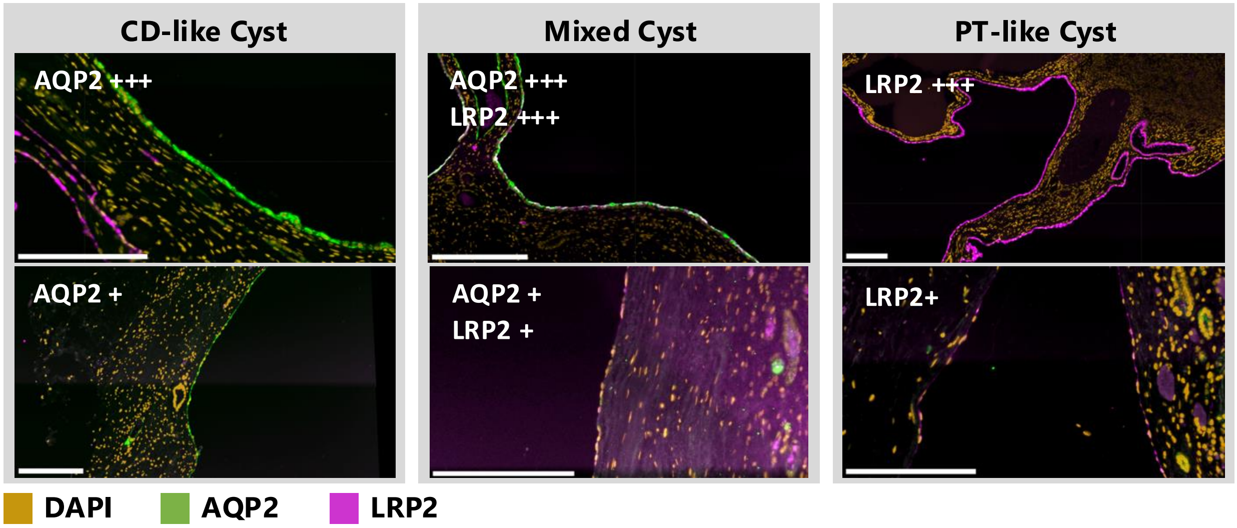


**Special Cases:**

AQP2+++ / LRP2 (+) → CD-like Cyst
AQP2 + / LRP2 (+) → CD-like Cyst

LRP2 +++ / AQP2 (+) → PT-like Cyst
LRP2 + / AQP2 (+) → PT-like Cyst

Supplementary Figure 14

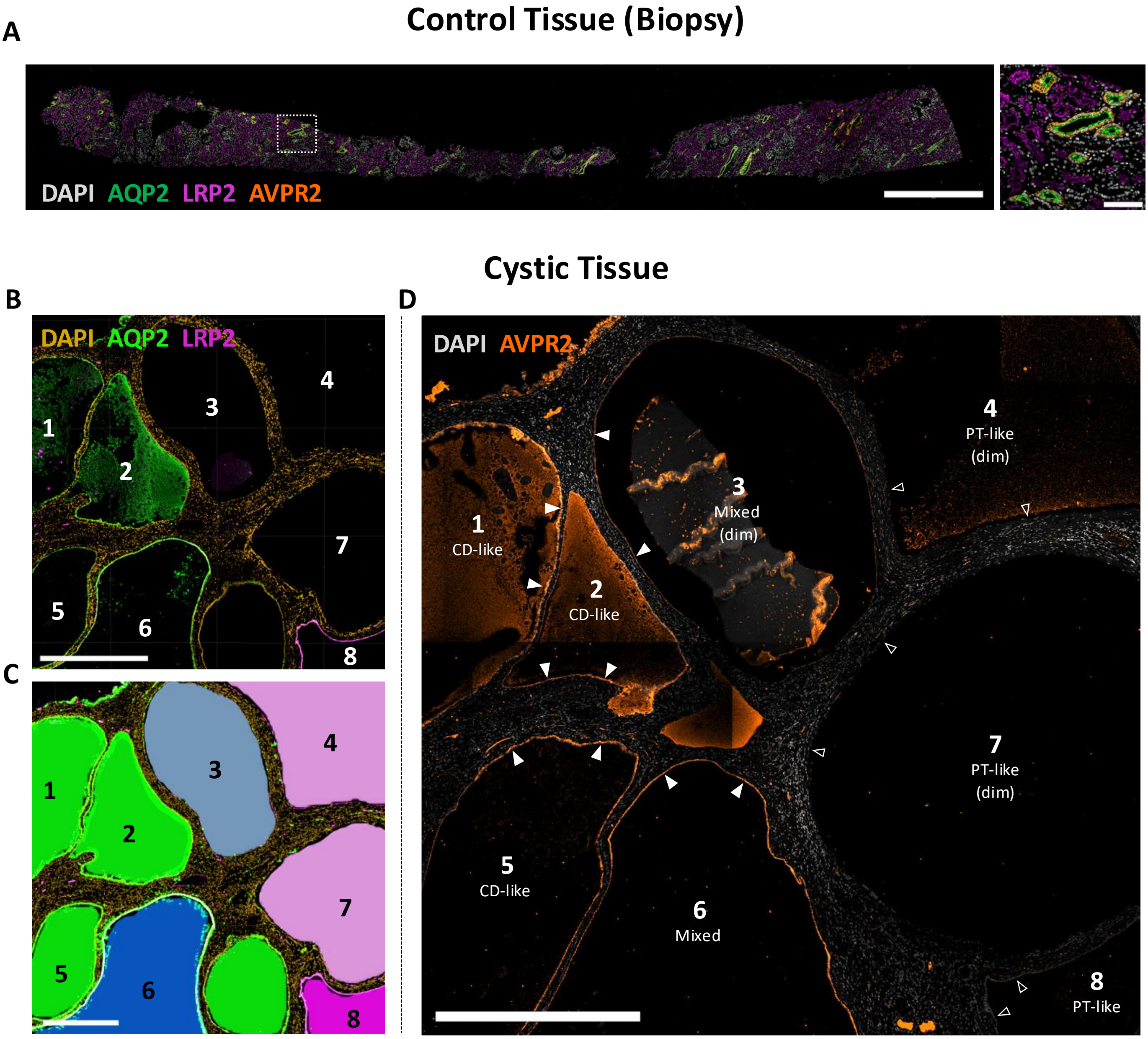

### Supplementary Figure 15

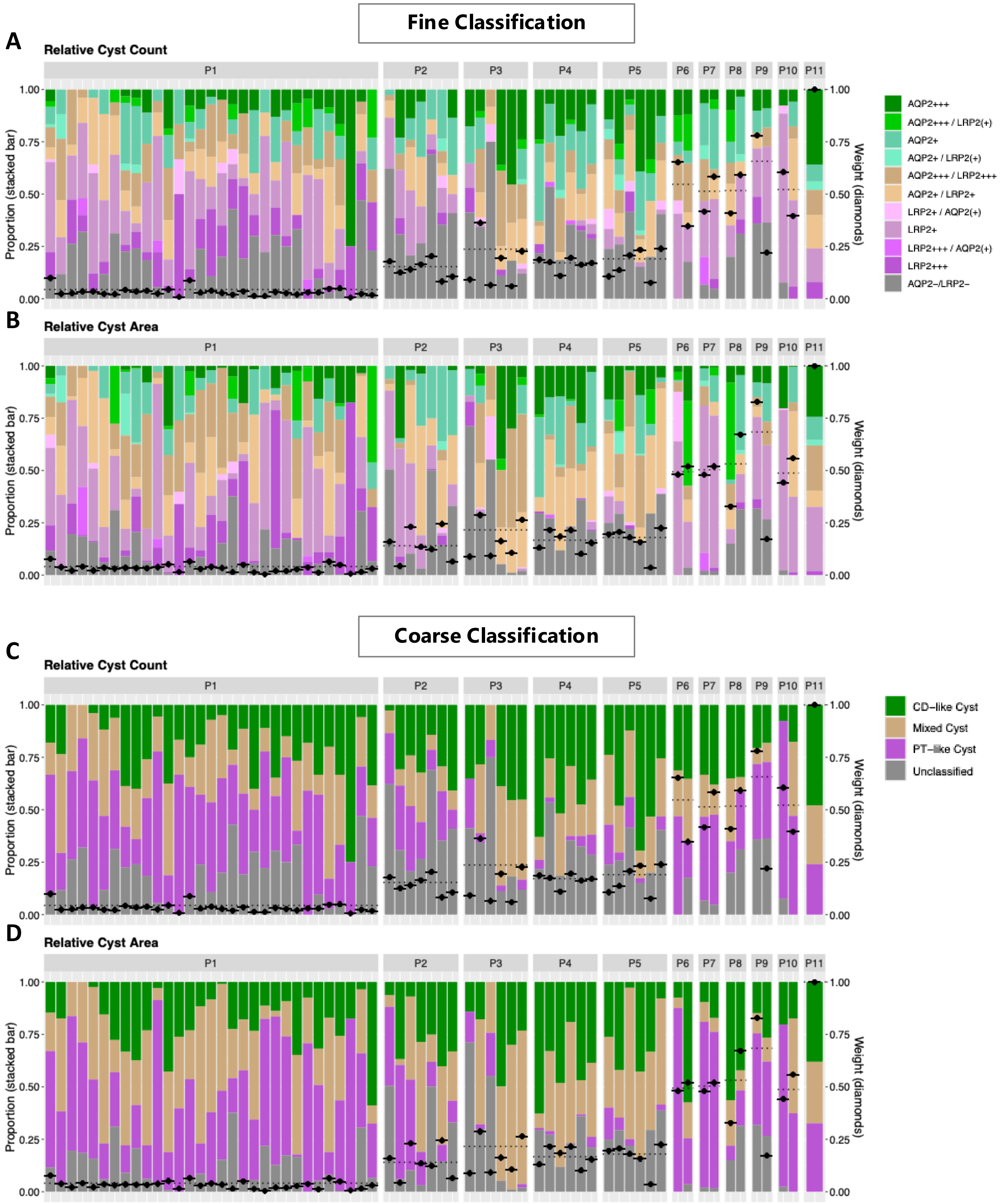

Supplementary Figure 16

**Fine Classification**

A

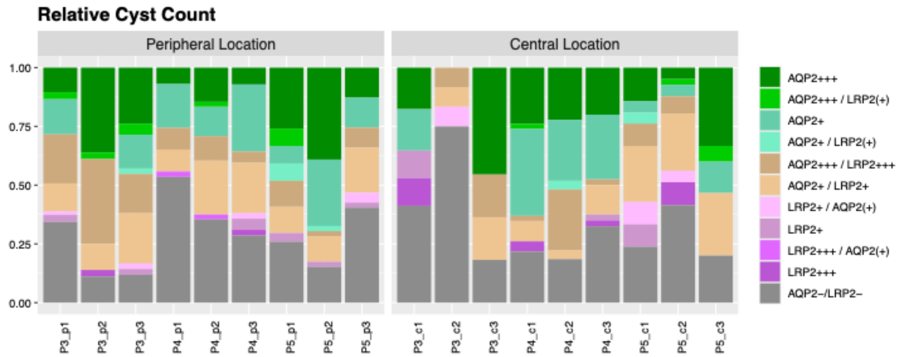


B

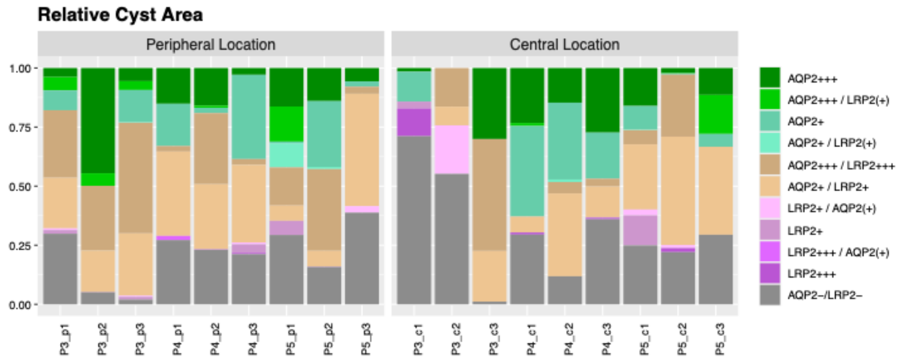


**Coarse Classification**

C

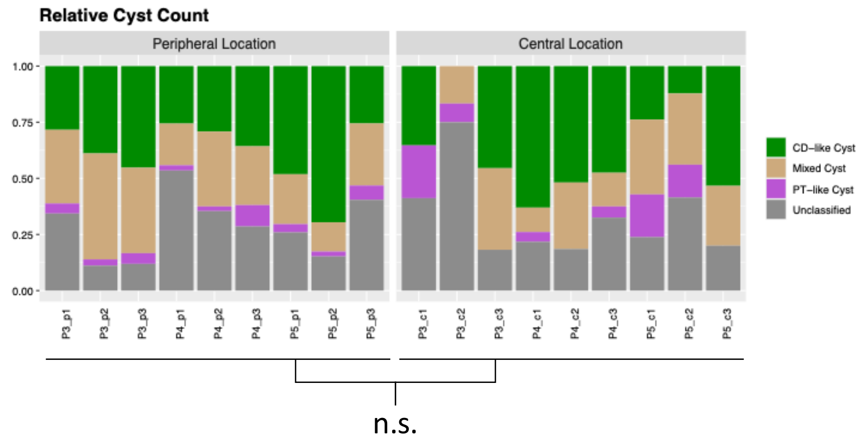


D

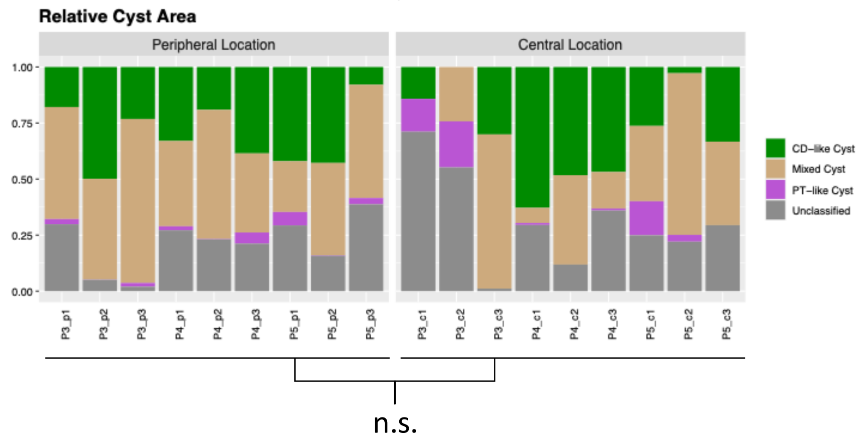

### Supplementary Figure 17

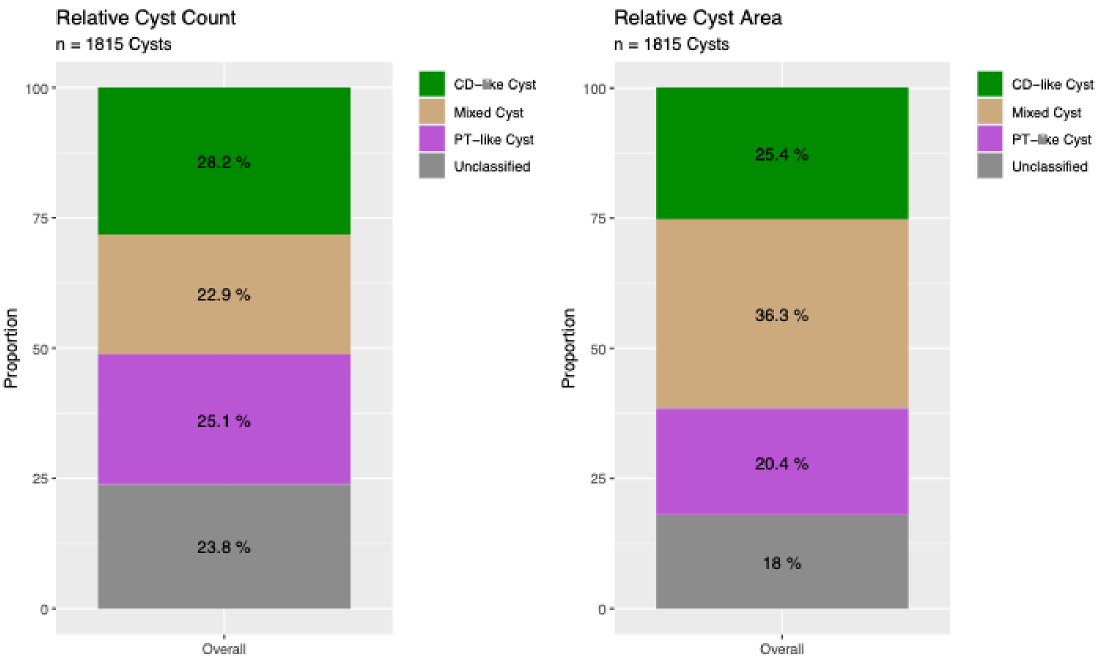

Supplementary Figure 18

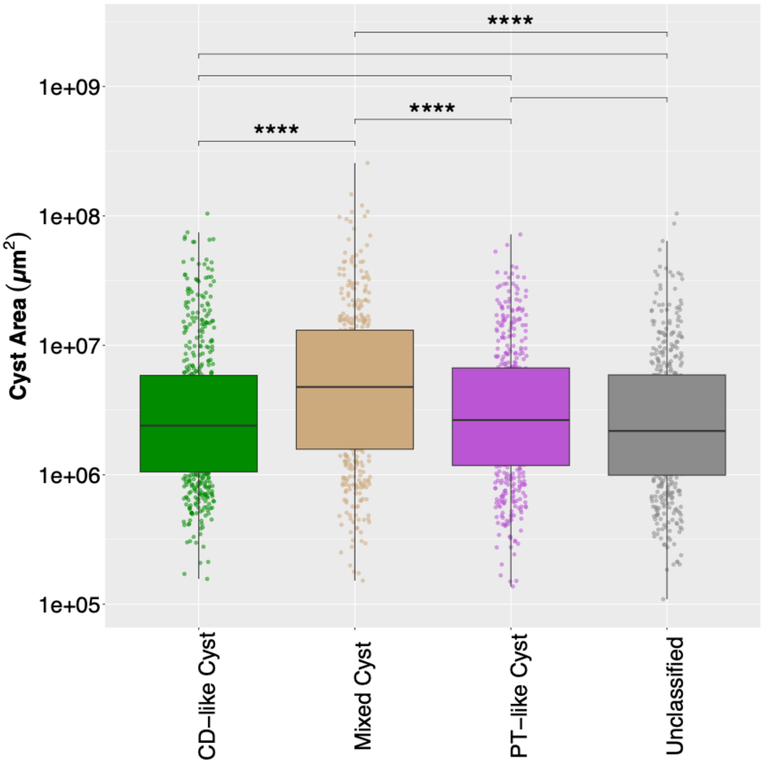

Supplementary Figure 19

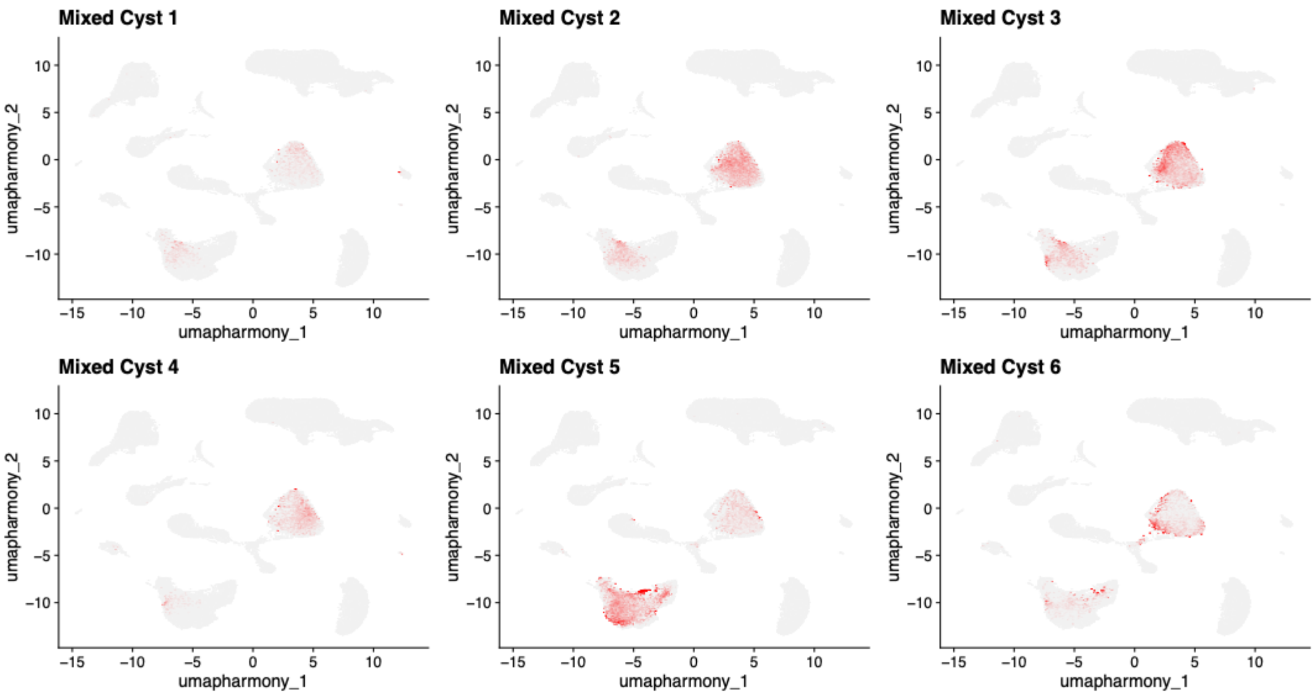
